# BCOR-Rearranged Sarcomas: *In Silico* Insights into Altered Domains and Reduced RAWUL-PUFD Binding

**DOI:** 10.1101/2024.05.21.595144

**Authors:** Kristóf Madarász, János András Mótyán, Yi-Che Chang Chien, Judit Bedekovics, Szilvia Lilla Csoma, Gábor Méhes, Attila Mokánszki

**Affiliations:** Department of Pathology, Faculty of Medicine, University of Debrecen, 4032 Debrecen, Hungary; Department of Biochemistry and Molecular Biology, Faculty of Medicine, University of Debrecen, 4032 Debrecen, Hungary

**Keywords:** BCOR, Fusion, PRC1, PUFD, Sarcoma

## Abstract

*BCOR* (BCL-6 corepressor)-rearranged small round cell sarcoma (BRS) is a rare soft tissue tumor, mostly featuring the *BCOR*::*CCNB3* fusion, with other fusions like *BCOR*::*MAML3*, *BCOR*::*CLGN*, *ZC3H7B*::*BCOR*, *KMT2D*::*BCOR*, *CIITA*::*BCOR*, and *RTL9*-*BCOR* also reported. BCOR, a Polycomb Repressive Complex 1 (PRC1) component, influences histone modifications. It dimerizes with Polycomb group RING finger homolog (PCGF1) via its PCGF ubiquitin-like fold discriminator (PUFD) domain interacting with PCGF1’s RING finger and WD40-associated ubiquitin-like (RAWUL) domain. We used various *in silico* tools to explore the impact of fusion events on BCOR’s functionality and RAWUL-PUFD dimer binding affinity. Changes were found in the domain landscapes, physicochemical properties, GO terms and significant increases in the disordered regions within the PUFD domain of the fusion proteins. Structural predictions indicated modified intermolecular contacts (ICs) and a significant reduction in binding affinity in fusion protein RAWUL-PUFD dimers. These findings align with expression data showing PRC1-regulated gene upregulation in BRS, likely due to reduced RAWUL-PUFD binding affinity, impacting dimer formation and PRC1 assembly. Our findings enhance the understanding of BRS oncogenesis and identify potential therapeutic targets.

## Introduction

BRS is a rare soft tissue tumor occurring in the bones of young patients. Based on the 5^th^ edition of WHO Classification of soft tissue and bone tumors, BRS belongs to the third distinct subset of "undifferentiated small round cell sarcomas of bone and soft tissue" (Sbaraglia *et al*, 2020) with histological features, immune profile, and expression signatures, that differ from round cell sarcomas with *EWSR1* gene fusion with non-ETS family members and from Capicua transcriptional repressor (*CIC)*-rearranged sarcomas (Kao *et al*, 2018).

The *BCOR* gene is located in the Xp11.4 chromosomal region and mediates apoptotic and oncogenic activities of cells through *BCL6*-regulated transcriptional repression *via* epigenetic signaling mechanisms (Albagli *et al*, 1999; Huynh *et al*, 2000; Pagan *et al*, 2007). BCOR (Uniprot ID: Q6W2J9) binds to PCGF1 of PRC1 through its PUFD domain, which facilitates epigenetic modification of histones by the addition of a ubiquitin moiety to histone H2A at Lys119 (H2AK119) (Junco *et al*, 2013; Blackledge *et al*, 2015). The BCOR PUFD and the RAWUL domain interact in a hierarchical mode of assembly. Upon binding of PCGF1, the BCOR PUFD termini become structurally ordered, enabling stable association with KDM2B. This interaction is critical for the assembly of the noncanonical PRC1 variant known as PRC1.1 (Blackledge *et al*, 2014; Wong *et al*, 2020). The interaction between RAWUL and PUFD depends on the β-sheet and loop interaction surfaces for selective binding of their respective partners (Junco *et al*, 2013). PRC1 silences several genes, including the HOX group of genes (Blackledge *et al*, 2015; Chittock *et al*, 2017). It inhibits the mesoderm and endoderm specification genes and assumes an integral part in regulating the primed pluripotent state. Notably, BCOR seems to be non-essential for mouse embryonic stem cell (mESC) pluripotency similar to other recently reported human ESC (hESC) regulators (Wamstad *et al*, 2008). Recruitment of BCOR to the target sites is primarily mediated by its C terminus, which plays a necessary and sufficient role in this process. The PUFD region is likely to play a key role in targeting BCOR to PRC1.1. In addition, the N terminus of BCOR contributes to transcriptional repression independently of the C terminus (Wang *et al*, 2018).

Internal tandem duplications (ITDs) and translocations leading to gene fusions are the most frequently occurring aberrations of *BCOR.* The G2/mitotic-specific cyclin-B3 (*CCNB3*) was identified as the most prevalent fusion partner in BRS (Pierron *et al*, 2012; Kao *et al*, 2016). Following the increasing use of molecular diagnostics methods, especially next-generation sequencing (NGS), Spacht *et al*. reported two novel fusions in 2016, namely *BCOR*::*MAML3* (mastermind-like protein 3) and *ZC3H7B*::*BCOR* (ZC3H7B: zinc finger CCCH domain-containing protein 7B) (Specht *et al*, 2016). Subsequently, Kao *et al*. discovered the *KMT2D*::*BCOR* fusion (KMT2D: histone-lysine N-methyltransferase 2D) in 2018 (Kao *et al*, 2018), while Yoshida *et al*. detected *CIITA*::*BCOR* (CIITA: MHC class II transactivator) and *ZC3H7B*::*BCOR* fusions in 2020 (Yoshida *et al*, 2020). More recently, in 2022, Vassella *et al*. identified an *RTL9*::*BCOR* gene fusion (*RTL9:* retrotransposon gag-like protein 9; the protein is also referred to as retrotransposon gag domain-containing protein 1, *RGAG1*). As compared to the other fusions described above, this fusion protein uniquely contains a non-canonical BCOR that lacks its 1168-1201 region and thus corresponds to isoform 2 of BCOR (BCOR-2) (Vasella *et al*, 2022). An additional fusion was reported in 2023, Chang Chien *et al*. described the *BCOR*::*CLGN* (calmegin) fusion in BRS (Chang Chien *et al*, 2023).

In this study, we aimed (i) to collect the currently available literature data about BRS, including cDNA and amino acid sequences of the fusion genes based on NGS and Real-Time-PCR (RT-PCR) data, (ii) to use *in silico* approaches for the analysis of chimeric sequences as well as structures, and (iii) to build the 3D structures of the chimeric proteins, to reveal the oncogenic mechanism of *BCOR* fusion small round cell sarcomas.

## Results

### Sequence analysis

After collecting the nucleotide sequences (**Appendix Text S1**), we performed the translation of the sequences *in silico*, followed by the analysis of the fusion proteins’ sequences, including alignment of the sequences of the fusion proteins with those of the wild-type proteins (by using Blastp tool) and determination of the residual lengths of the partner proteins.

The seven studied fusion proteins were classified into two groups according to the position of the wild-type BCOR (wBCOR) within the fusion protein. In three of the seven cases, the wBCOR protein was located at the N terminus, while in the other four cases at the C terminus of the fusion proteins. These groups were referred to as BCOR^NT^ and BCOR^CT^, respectively.

The BCOR was present at the N terminus of the BCOR::CCNB3, BCOR::CLGN and BCOR::MAML3 fusion proteins **Fig. 1**. The BCOR::CCNB3 (UniProt ID: H9A532; 3038 AA) and BCOR::CLGN (2070 AA) were the only fusion proteins that encompass the full-length wBCOR (1755 AA). In these cases, the fusion caused the loss of a short N-terminal region of CCNB3 (<10% of the full-length protein sequence), while the truncation was more remarkable (∼50%) in CLGN where 1-295 of the 610 residues were deleted upon the fusion, as previously reported (Chang Chien *et al*, 2023). The BCOR::MAML3 fusion proteins contained the nearly full-length wBCOR, the 1-1751 residues were present and only the 1751-1755 C-terminal residues were missing. The 1752-2734 region of the fusion protein corresponded to the MAML3 protein having a truncated N terminus, and 1-156 of the 1138 residue long protein was missing. The amino acid at position 1752 was formed by guanine, alanine of *BCOR,* and by guanine base of *MAML3*.

**Figure 1.**
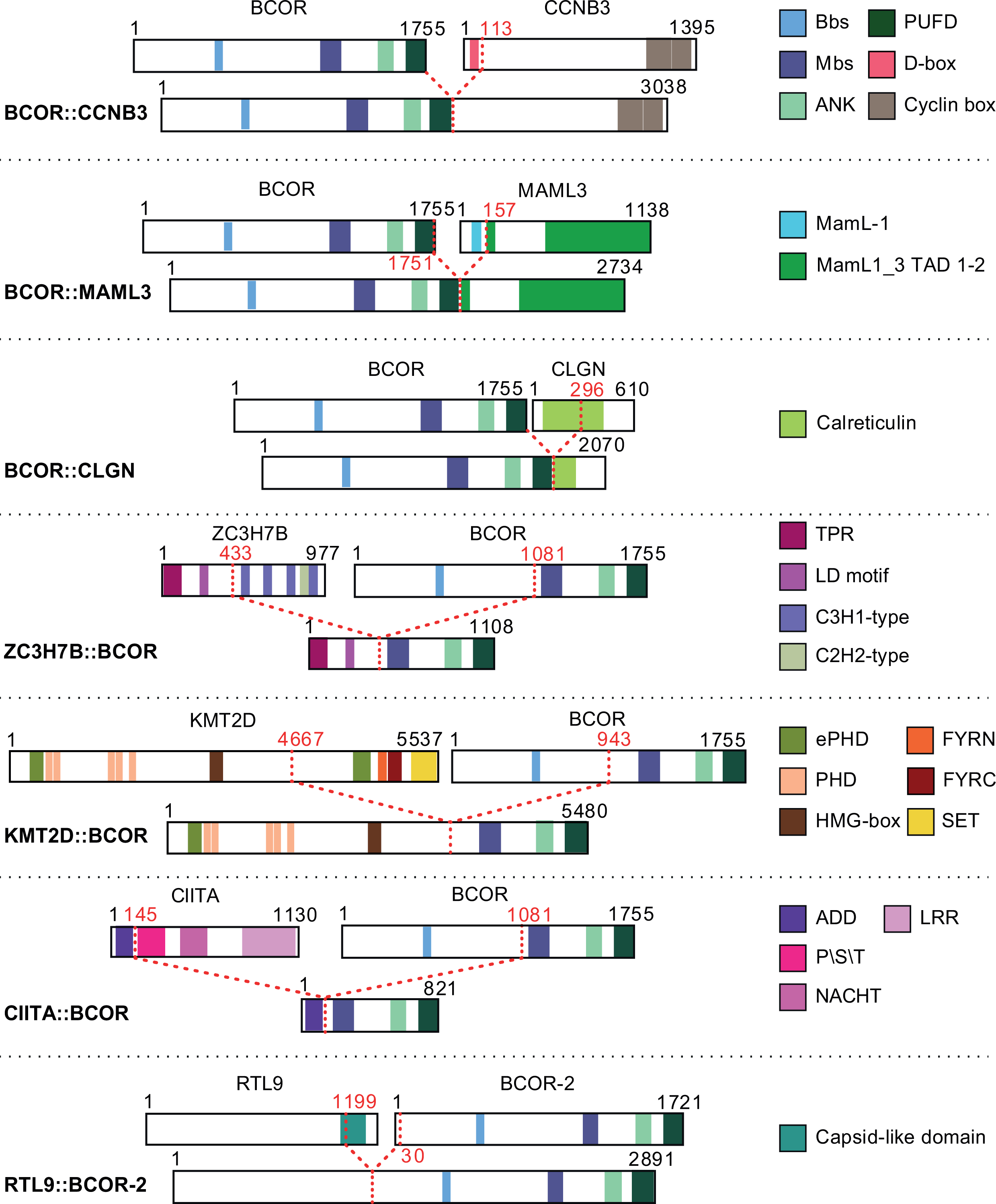
Domains and functional sites of known fusion proteins from *BCOR*-rearranged sarcomas. As a result of the fusion events, the new fusion protein has lost several regions, both short and long, along with the domains and functional sites when compared to the wild-type fusion partners. Schematic representation is shown for each fusion partner and fusion protein, in the order corresponding to their N-or C-terminal positions in the fusion proteins. The red dotted lines and labels indicate the sites of the breakpoints at the protein level. The lengths of KMT2D and KMT2D::BCOR proteins, except for the domains, are not representative due to the large size of these two proteins, all other proteins are represented proportionally. Bbs: BCL6 biding site, Mbs: MLLT biding site, ANK: ankyrin repeats; PUFD: PCGF Ubiquitin-like fold discriminator; D-box: destruction box, MamL-1: Neurogenic mastermind-like, N-terminal domain, Maml1_3 TAD 1-2: Mastermind-like 1/3: transactivation domain; TPR: Tetratricopeptide repeats, LD motif: leucine-aspartate repeat motif, C3H1-type: CCCH type, C2H2-type: CCHH type; ePHD: extended plant homeodomain, PHD: plant homeodomain, HMG-box: high mobility group -box, FYRN: FY-rich domain N-terminal region, FYRC: FY-rich domain C-terminal region, SET: pre-SET and post-SET region; ADD: acetyltransferase domain, P\S\T: Proline-serine-threonine rich domain, LRR: Leucine-rich repeat ribonuclease inhibitor; BCOR-2: BCOR isoform 2.

In the case of BCOR^CT^ group, the fusion caused a more remarkable shortening of the BCOR protein sequence, except RTL9::BCOR-2. In the other three cases, there was a 54-62% decrease in the length of the wBCOR sequence (**Fig. 1**, **Table 1**). The length of the ZC3H7B::BCOR fusion protein was 1108 AA, its 1-433 region corresponds to the truncated ZC3H7B protein while the 434-1108 region to the 1081-1755 residues of wBCOR. In the case of *KMT2D*::*BCOR*, the fusion resulted in a 5480 AA long fusion protein. It represented the 1-4667 and 943-1755 residues of KMT2D and wBCOR, respectively, the 16% and 54% of the respective full-length KMT2D and BCOR sequences were missing from the fusion proteins.

**Table 1.** Fusion events and breakpoints (BP) in *BCOR*-rearranged sarcomas. AA: amino acids, wt: wild type. The fusion of the *RTL9* and *BCOR* genes results in the RTL9::BCOR-2 fusion protein. BCOR-2 refers to isoform 2 of BCOR which consists of 1721 residues. In the case of the other fusions, the BCOR corresponds to the canonical form containing 1755 residues. A schematic representation of the fusion proteins is shown in **Fig. 1**. The lengths are shown for the fusion partners if they are part of the fusion protein (partner 1 or 2) or not (wt).

| Index | Gene information |  |  |  |  | Protein information |  |  |
| --- | --- | --- | --- | --- | --- | --- | --- | --- |
|  | Fusions | Gene 5' BP | Gene 3' BP | Chromosomal partners | Exonic BP 5' - 3' | Partner 1 / wt length (AA) | Partner 2 / wt length (AA) | Fusion protein length (AA) |
| 1 | *BCOR::CCNB3* | 39 911 366 | 50 051 504 | chr X - chr X | 15. - 5. | 1755 / 1755 | 1283 / 1395 | 3038 |
| 2 | *BCOR::MAML3* | 39 911 374 | 141 074 014 | chr X - chr 4 | 15. - 1. | 1751 / 1755 | 982 / 1138 | 2734 |
| 3 | *BCOR::CLGN* | 39 911 366 | 141 317 359 | chr X - chr 4 | 15. - 9. | 1755 / 1755 | 316 / 610 | 2070 |
| 4 | *ZC3H7B::BCOR* | 41 738 632 | 39 923 852 | chr X - chr X | 12. - 7. | 433 / 977 | 675 / 1755 | 1108 |
| 5 | *KMT2D::BCOR* | 49 424 063 | 39 931 774 | chr 12 - chr X | 42. - 4. | 4667 / 5537 | 813 / 1755 | 5480 |
| 6 | *CIITA::BCOR* | 10 992 859 | 39 923 852 | chr 16 - chr X | 5. - 7. | 145 / 1130 | 675 / 1755 | 821 |
| 7 | *RTL9::BCOR* | 109 697 442 | 39 937 097 | chr X - chr X | 2. - 3. | 1199 / 1388 | 1692 / 1721 | 2891 |

The most remarkable sequence truncation was observed for the fusion partners in the case of the CIITA::BCOR, which was the shortest of them (containing 821 AA). The fusion protein contained only 13% of the CIITA and 38% of the wBCOR sequences, representing the 1-145 N-terminal and 1081-1755 C-terminal regions of the wild-type proteins, respectively. The triplet coding for the Arg146 residue was constituted by a cytosine of the *CIITA* and guanine and cytosine of the *BCOR* gene.

The RTL9::BCOR-2 fusion protein – consisting of 2891 residues – contained the RTL9 protein truncated at its C terminus (the 1200-1388 region is missing) as well as the nearly full-length BCOR-2 (UniProt ID: Q6W2J9-2) which lacked only its 1-29 C-terminal residues in the fusion protein.

### Physicochemical characteristics of fusion proteins

The physicochemical properties of the fusion proteins were calculated by using the ProtParam web server tool (**Fig. 2**). In all instances, we observed alterations in the physicochemical properties of the fusion proteins as compared to the wild-type proteins (**Table 2**).

**Figure 2.**
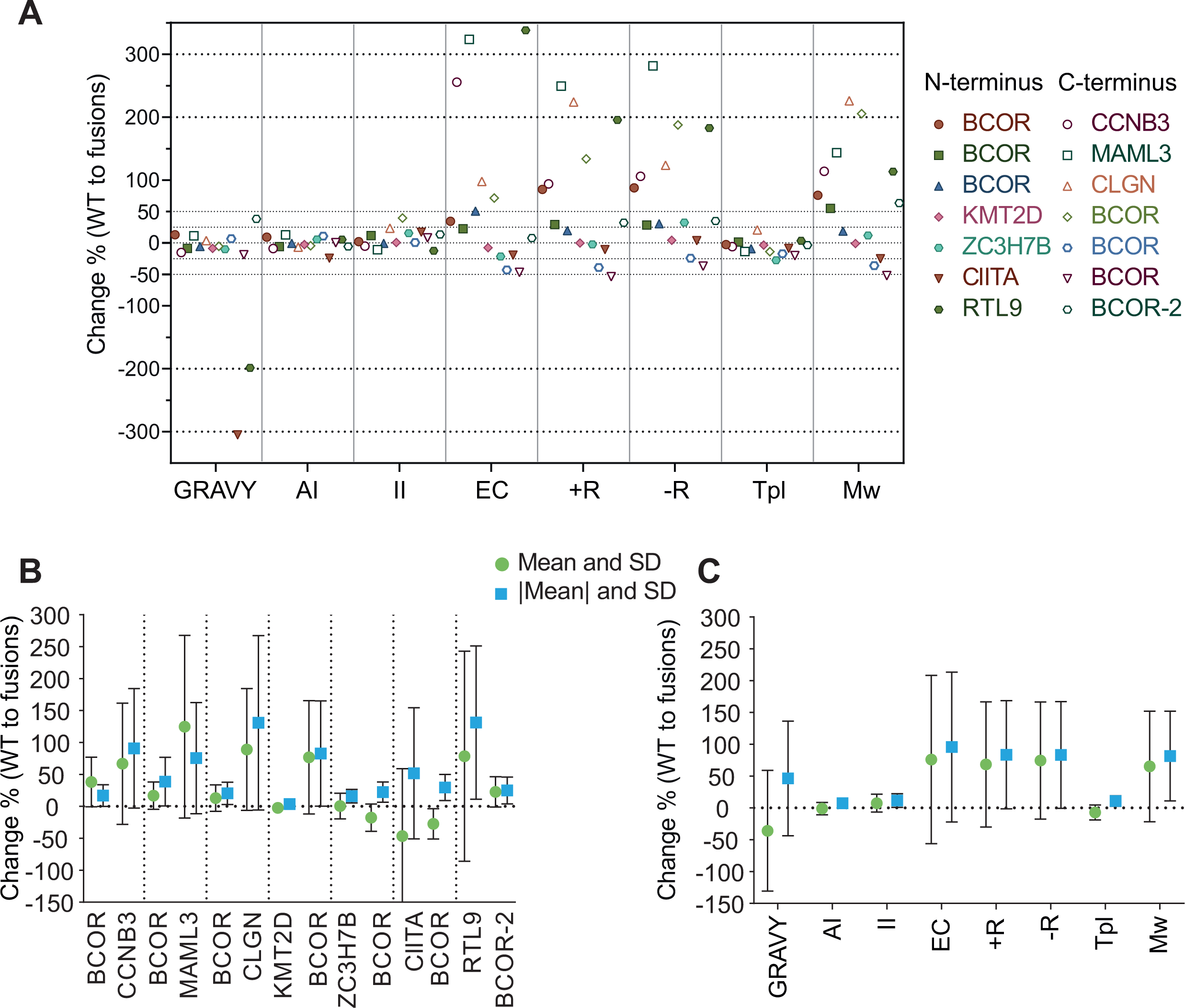
Changes in the fusion proteins’ physicochemical properties as compared to wild-type partner proteins. **(A)** Differences of eight investigated features (%) between wild-type and fusion proteins. GRAVY: grand average hydropathy, AI: aliphatic index, II: instability index, EC: extinction coefficient at 280 nm, +R: the number of positively charged residues, −R: the number of negatively charged residues, TpI: the theoretical isoelectric point, Mw: molecular weight, WT: wild-type. **(B)** The mean and standard deviation of the percentage changes in proteins. **(C)** The mean and standard deviation of the percentage changes in eight attributes. |Mean|: mean of absolute values, Error bars show standard deviation (SD).

**Table 2.** The physicochemical parameters were calculated for the wild-type and the fusion proteins. Partner’s average: the mean of the values for each pair of proteins involved in the fusion. Partner’s average represents the mean of the values for each pair of proteins involved in the fusion, while Fusion’s average represents the mean of the fusion protein’s value. GRAVY: grand average hydropathy, AI: aliphatic index, II: instability index, EC: extinction coefficient at 280 nm, +R: the number of positively charged residues, −R: the number of negatively charged residues, TpI: the theoretical isoelectric point, Mw: molecular weight, Sl: sequence length, AA: amino-acids, BCOR-2: Isoform 2 of BCOR.

|  | GRAY | AI | II | EC | + R | + R (%) | -R | -R (%) | TpI | Mw (Da) | SI (AA) |
| --- | --- | --- | --- | --- | --- | --- | --- | --- | --- | --- | --- |
| BCOR | -0.668 | 68.38 | 55.26 | 160115 | 203 | 8,65 | 226 | 7,77 | 6.06 | 192188.64 | 1755 |
| CCNB3 | -0.504 | 82.01 | 59.29 | 60500 | 194 | 7,19 | 206 | 6,77 | 6.28 | 157916.13 | 1395 |
| BCOR::CCNB3 | -0.581 | 74.72 | 56.34 | 215115 | 376 | 8,08 | 424 | 7,17 | 5.91 | 337723.51 | 3038 |
| MAML3 | -0.817 | 57.01 | 69.23 | 46340 | 75 | 15,17 | 76 | 14,97 | 7.12 | 122292.88 | 1138 |
| BCOR::MAML3 | -0.725 | 64.3 | 61.71 | 196235 | 262 | 10,44 | 290 | 9,43 | 6.15 | 297852.28 | 2734 |
| CLGN | -0.734 | 72.74 | 44.52 | 121850 | 75 | 8,13 | 132 | 4,62 | 4.57 | 70038.6 | 610 |
| BCOR::CLGN | -0.706 | 67.92 | 54.94 | 240930 | 243 | 8,52 | 295 | 7,02 | 5.51 | 228230.9 | 2070 |
| KMT2D | -0.648 | 67.03 | 76.61 | 296935 | 475 | 11,66 | 623 | 8,89 | 5.4 | 593389.41 | 5537 |
| KMT2D::BCOR | -0.704 | 65.27 | 77.16 | 274350 | 474 | 11,56 | 650 | 8,43 | 5.22 | 587199.74 | 5480 |
| ZC3H7B | -0.567 | 71.72 | 48.26 | 116710 | 127 | 7,69 | 129 | 7,57 | 6.94 | 109858.00 | 977 |
| ZC3H7B::BCOR | -0.624 | 75.73 | 55.68 | 91565 | 124 | 8,94 | 171 | 6,48 | 5.02 | 122865.76 | 1108 |
| CIITA | -0.196 | 90.72 | 50.97 | 105780 | 105 | 10,76 | 138 | 8,19 | 5.30 | 123513.99 | 1130 |
| CIITA::BCOR | -0.794 | 68.68 | 59.75 | 85480 | 94 | 8,73 | 143 | 5,74 | 4.84 | 92428.75 | 821 |
| BCOR-2 | -0.661 | 68.43 | 55.38 | 153125 | 201 | 8,56 | 218 | 7,89 | 6.22 | 188202.33 | 1721 |
| RTL9 | -0.137 | 61.27 | 71.76 | 37650 | 90 | 15,42 | 104 | 13,35 | 5.81 | 144279.61 | 1388 |
| RTL9::BCOR-2 | -0.409 | 64.52 | 62.82 | 164960 | 266 | 10,87 | 294 | 9,83 | 6.01 | 307716.11 | 2891 |
| Partner's average | -0,591 | 70.09 | 57.68 | 135684 | 183 | 9.75 | 213 | 8.49 | 6.00 | 190187.34 | 1739 |
| Fusion's average | -0,649 | 68.73 | 61.20 | 181234 | 263 | 9.59 | 324 | 7.73 | 5.52 | 282002.44 | 2592 |

The mean alteration across all cases was 30.90%, and the mean alteration of absolute values was 46.64% (|mean|). The percentages of +R(%) and -R(%) which compare the charged amino acids to the whole protein length were 10.02% and 8.38% respectively, with standard deviations of 2.48 and 2.63. Two main groups were established based on the localization of the BCOR in the fusion proteins. For the BCOR^NT^ group, the mean change was 58.03% and the absolute mean was 55.25%, based on the ProtParam attributes. In the case of the BCOR^CT^ proteins, the mean change was 10.54% and 40.18%, respectively (**Fig. 2**, **Table 2**).

The MAML3 protein exhibited the most substantial mean change of 124.69%, which can be explained by its large truncation due to the fusion event. In contrast, the smallest mean difference (0.47%) and the smallest |mean| change (3.56%) were observed for ZC3H7B and KMT2D, respectively. The mean change of the GRAVY values was -35.95% (**Fig. 2C**), with the smallest and largest differences observed for CLGN (3.81%) and CIITA (-305.10%). The mean change in AI values was negligible, although the CIITA showed a 24.29% decrease, resulting in a decrease in thermostability. At the II, the mean alterations were more noticeable, with the greatest change found in the case of BCOR protein in KMT2D::BCOR fusion, which increased by 39.63%, indicating a decrease in protein thermostability. The mean change in +R and -R was 68.3% to 74.43%, with the highest being 249.33% and 281.58%, both in MAML3, due to its drastic truncation due to the fusion event. The average TpI decreased by 7.41%. however, none of the fusion proteins exceeded the pH range of 4.5-10, indicating nuclear localization (Schwartz *et al*, 2001).

The CIITA protein’s AI and GRAVY values decreased by -305.10% and -24.29%, respectively, indicating a decrease in protein thermostability and an increase in hydrophilicity. It is noteworthy that the BCOR protein in the KMT2D::BCOR fusion showed an increase in II, despite already having a high II value and a longer protein length than BCOR. This resulted in a considerable decrease in the stability of the BCOR partner, with little change in the stability of KMT2D (**Fig. 2C**).

Overall, considerably lower SD values were observed for the AI, II, and TpI values than for the other parameters (**Fig. 2C**). The highly similar aliphatic index values (AI) imply similar thermal stability for the fusion proteins. The instability indexes are also similar, the highest value was calculated for KMT2D::BCOR, indicating the lowest relative stability for this protein. The comparable averages that were calculated for the wild-type partners and the fusion proteins indicate only moderate change for these parameters.

Based on the comparison of the mean values of the partner proteins and the fusion proteins, the Gravy, AI, -R(%) and +R(%) values of the partner proteins showed a decrease, while II was increased. This led to an increase in hydrophobicity, a decrease in thermostability, a lower ratio of charged amino acids to total protein length, and a higher probability of protein degradation or denaturation.

### Protein domains and structural changes

The CDD, the UniProt, and the InterPro databases were utilized to map the domains and interaction sites for the standalone wild-type and the fusion proteins.

Two of the four relevant regions of the BCOR protein, namely the Ankyrin (ANK) repeats and the PUFD domain were present in all the novel proteins after the fusion events (**Fig. 1**). Regarding the remaining two sites, the BCL6 binding site (Bbs) and the MLLT3 binding site (Mbs), it was found that the Mbs was present in all fusion proteins. However, it should be noted that the Mbs of BCOR-2 - in the RTL9::BCOR-2, as well - consists of 89 residues while that of the canonical BCOR contains 123 residues (the Mbs sequences differ only in their length and show 100% sequence identity). The Bbs were observed in all three BCOR^NT^ proteins but were present in the RTL9::BCOR-2 of the BCOR^CT^ group.

When considering other fusion partners, a mosaic picture arises regarding the functional domains kept and lost.

The fusion resulted in the loss of a short N-terminal region of CCNB3 containing its destruction box (D-box), whereas the BCOR::CCNB3 protein contained both C-terminal cyclin boxes of CCNB3 (**Fig. 5A**).

The BCOR::MAML3 fusion protein does not contain the N-terminal region of MAML3 that encompasses its neurogenic mastermind-like, N-terminal (MamL-1) domain as well as the first seven residues of MamL1/3 transactivation domain 1 (MamL1/3 TAD1). The other MAML3 regions including the long C-terminal MamL1/3 TAD2 domain are present in the fusion protein.

In the case of CLGN, the fusion resulted in the deletion of the majority of the calreticulin domain (232 of the 368 residues) which is responsible for calcium ion binding (Chang Chien *et al*, 2023).

The extended plant, the homeodomain (ePHD) zinc finger, the five PHDs, and the high mobility group (HMG) box domains of KMT2D were preserved, while the second ePHD, the F/Y-rich N terminus (FYRN), the F/Y-rich C terminus (FYRC) and the SET domain (including the post-SET domain) were absent from the fusion protein.

The ZC3H7B retained the LD motif and all three tetratricopeptide (TPR) repeats but its four C3H1-type zinc finger domains and a single C2H1-type zinc finger domain were missing from the ZC3H7B::BCOR protein.

Additionally, only the acetyltransferase domain of the CIITA protein remained intact in the case of the CIITA::BCOR fusion, whereas the CIITA’s P\S\T rich domain, NACHT domain, and leucine-rich repeats (LRRs) were lost.

Finally, the fusion event occurred at the 1199th position, affecting the 1169-1316 region of RTL9 near its C terminus. This region included both the N- and C-terminal subdomains (NTD and CTD) of its capsid-like domain. As a result of this fusion, almost the entire capsid-like domain was deleted (**Fig. 1**).

### Gene ontology

We conducted an InterPro search to acquire PANTHER GO terms for defining the functional attributes and to compare the biological processes, molecular functions, and cellular components of both the wild-type and fusion proteins. The results suggest that the GO terms associated with the BCOR protein, including negative regulation of transcription by RNA polymerase II (GO:0000122), transcription corepressor activity (GO:0003714), and nucleus (GO:0005634), were prevalent in the fusion proteins. As a result, the GO terms of the wild-type proteins were not assigned to almost all fusion partners except BCOR. Interestingly, the novel KMT2D::BCOR protein retained the GO terms of only the wild-type KMT2D but not those of the BCOR (**Table 3**). GO terms were not assigned to RTL9 and the RTL9::BCOR-2 proteins.

**Table 3.**
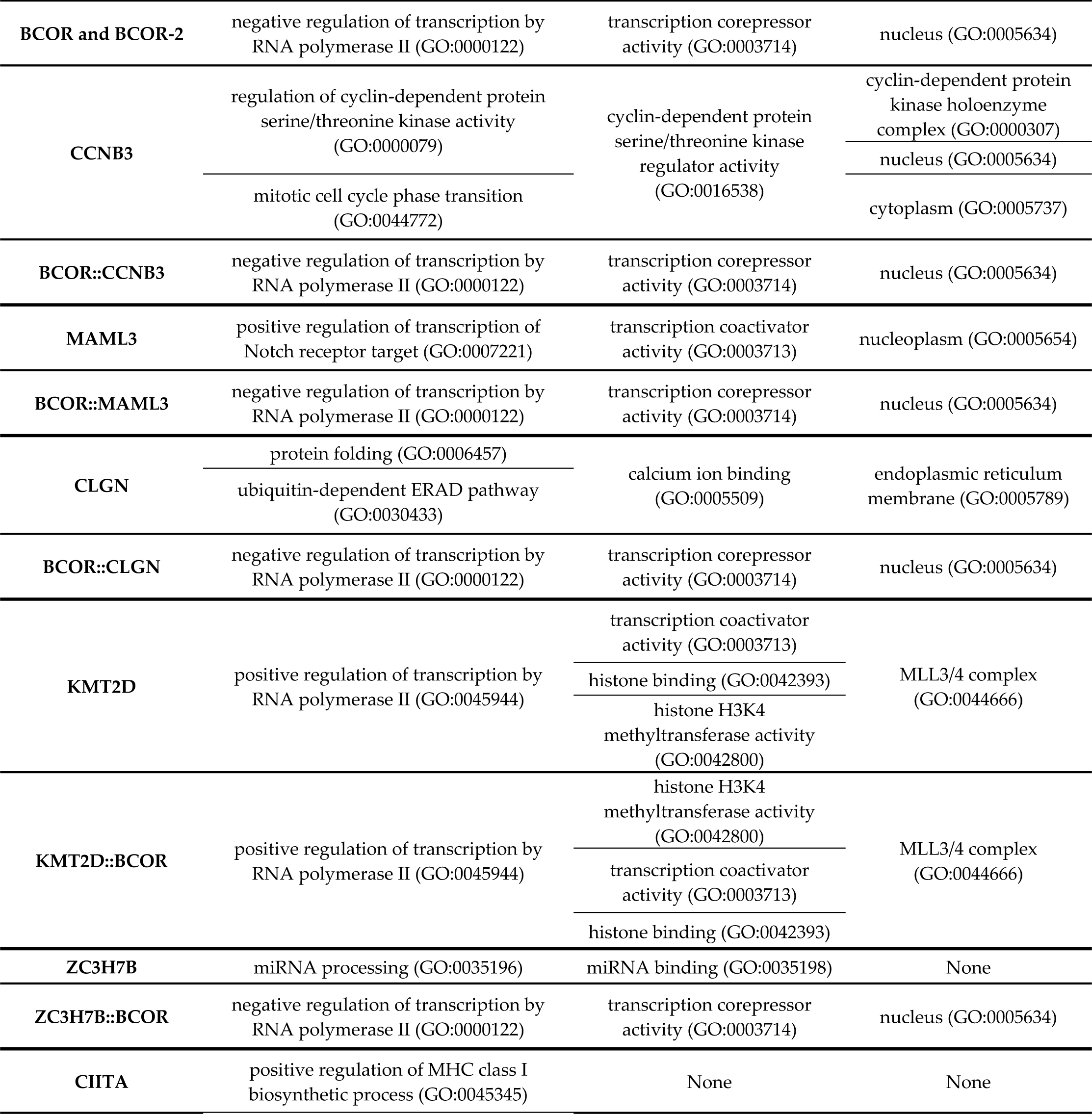
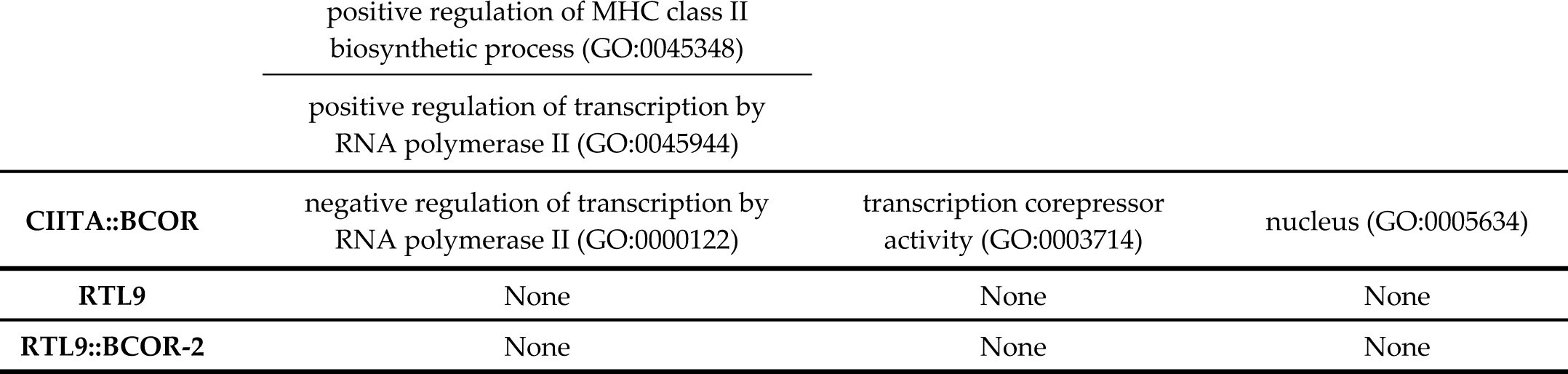
PANTHER Gene Ontology (GO) terms for partner proteins in their wild-type form and fusion pairs.

### Predicted intracellular localization and signal peptides

The SignalP 6.0 tool was utilized to identify standard secretory signal peptides (Sec/SPI), while the DeepLoc 2.0 tool was used to determine intracellular localization and calculate probability values for localization in the cytoplasm, nucleus, extracellular cell membrane, mitochondrion, plastid, endoplasmic reticulum (ER), lysosome, Golgi apparatus, and peroxisome. Additionally, the NetGPI-1.1 predictor was used to identify GPI-anchoring signals (**Table 4**).

**Table 4.**
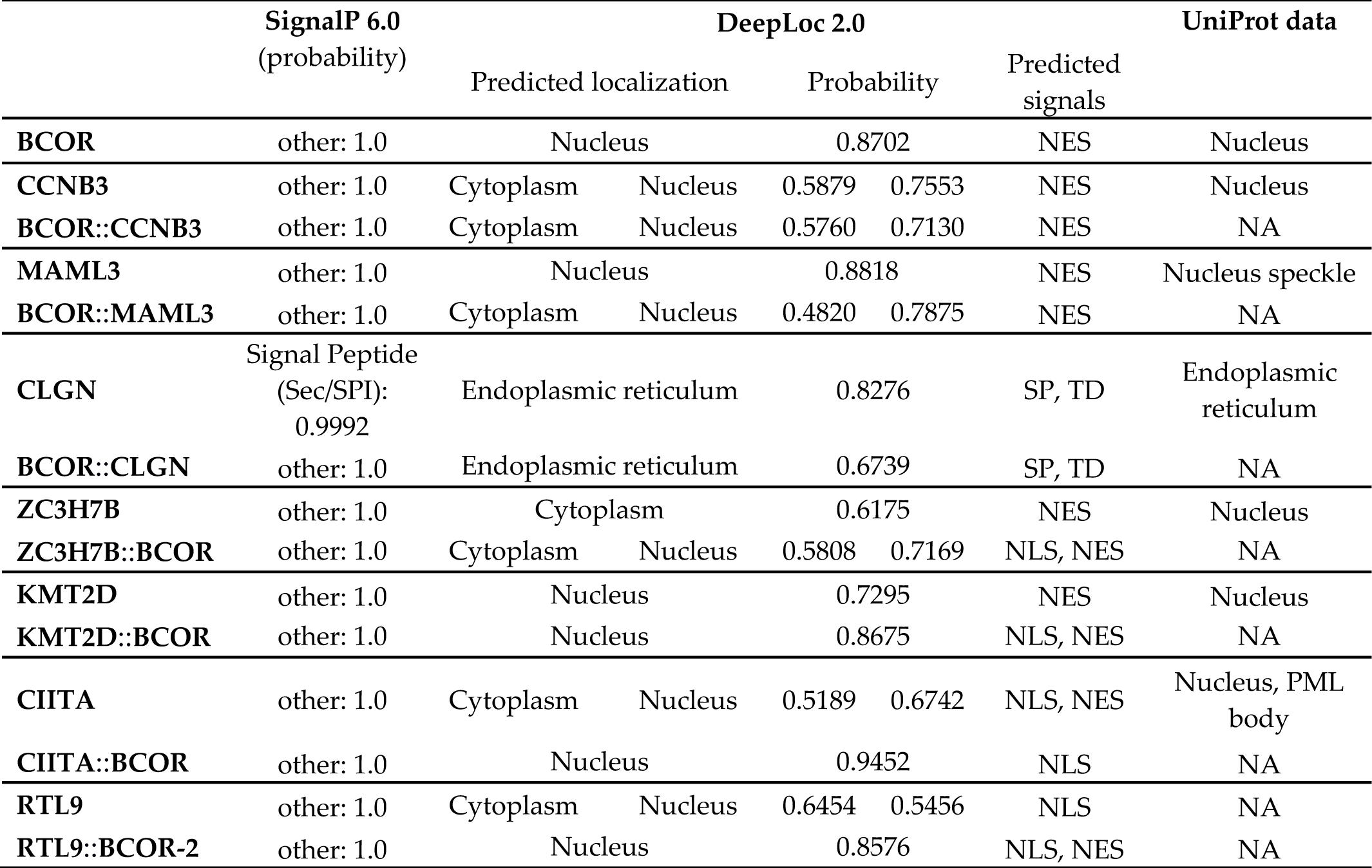
Intracellular localization and signals predicted for the wild-type and fusion proteins by using DeepLoc 2.0. NES. : nuclear export signal, **SP**: Sec signal peptide (Sec/SPI), **TD**: transmembrane domain, **NLS**: nuclear localization domain, **ER**: endoplasmic reticulum, **NA**: no data available, **PML**: Promyelocytic leukemia nuclear bodies, **other**: no signal peptides were found. For comparison, subcellular localization is indicated based on the UniProt database, if available. For DeepLoc predictions, only localizations reaching the threshold are shown.

According to the results of SignalP 6.0 analysis, only the CLGN protein contains a signal peptide, especially a secretory signal peptide with a probability of 99.92%, although, the N-terminal region of CLGN encompassing this sequence motif is missing from the BCOR::CLGN fusion.

The DeepLoc 2.0 prediction indicated that five of the eight wild-type proteins (BCOR, CCNB3, MAML3, KMT2D, and CIITA) were most likely located in the nucleus, while CLGN was predicted to be located in the endoplasmic reticulum, which was consistent with the data available in the UniProt database. Based on the data available in the Human Protein Atlas database, KMT2D may have plasma membrane and cytosol localization. The subcellular localization of ZC3H7B seems to be uncertain, as the UniProt database information implies its nuclear localization but - in agreement with the data of Human Protein Atlas - we predicted that it was located in the cytoplasm (score: 0.6175). Data for the RTL9 protein were not available in the UniProt database. The predicted score of cytoplasmic localization was 0.6454, while that of the nuclear localization was also similar (score: 0.5456). Most recent findings revealed that mouse RTL9 protein is intracellular and is highly expressed in microglial lysosomes in the neonatal brain (Ishino *et al*, 2023). All fusion proteins were predicted to be located in the nucleus except BCOR::CLGN protein which may be localized in the endoplasmic reticulum localization (predicted score: 0.6739).

DeepLoc 2.0 also predicted signals and found that all wild-type and fusion proteins, except for CLGN, BCOR::CLGN, CIITA::BCOR, and RTL9 proteins, contained a nuclear export signal (NES). The CLGN protein contained a signal peptide and a transmembrane domain, while BCOR::CLGN only contained a signal peptide. Nuclear localization signal (NLS) was predicted in the case of ZC3H7B::BCOR, KMT2D::BCOR, and RTL9::BCOR-2 next to the NES, while CIITA::BCOR and RTL9 protein only contained NLS according to DeepLoc 2.0 (**Table 4**). It should be mentioned that the prediction accuracy for NES is the lowest (<50%). This may explain its frequent occurrence among predictions.

Overall, the subcellular localization predicted by DeepLoc 2.0 agreed with the data available in UniProt and The Human Protein Atlas database in most cases (**Table 4**).

No GPI-anchoring signals were predicted by using the NetGPI-1.1 predictor, which implied that neither the wild-type proteins nor the fusion proteins contained a lipid anchor and were not cell-surface proteins.

### Prediction of Intrinsically Disordered Regions

We used the IUPred3 online tool to analyse the Intrinsically Disordered Regions (IDRs) of the proteins. Subsequently, we compared the region^ANK+linker^ and region^PUFD^ of the fusion proteins to the BCOR protein and in addition, the IUPred3 scores were also integrated into our statistical analyses by conducting Friedman’s tests on the values of each region separately (**Fig. 3**).

**Figure 3.**
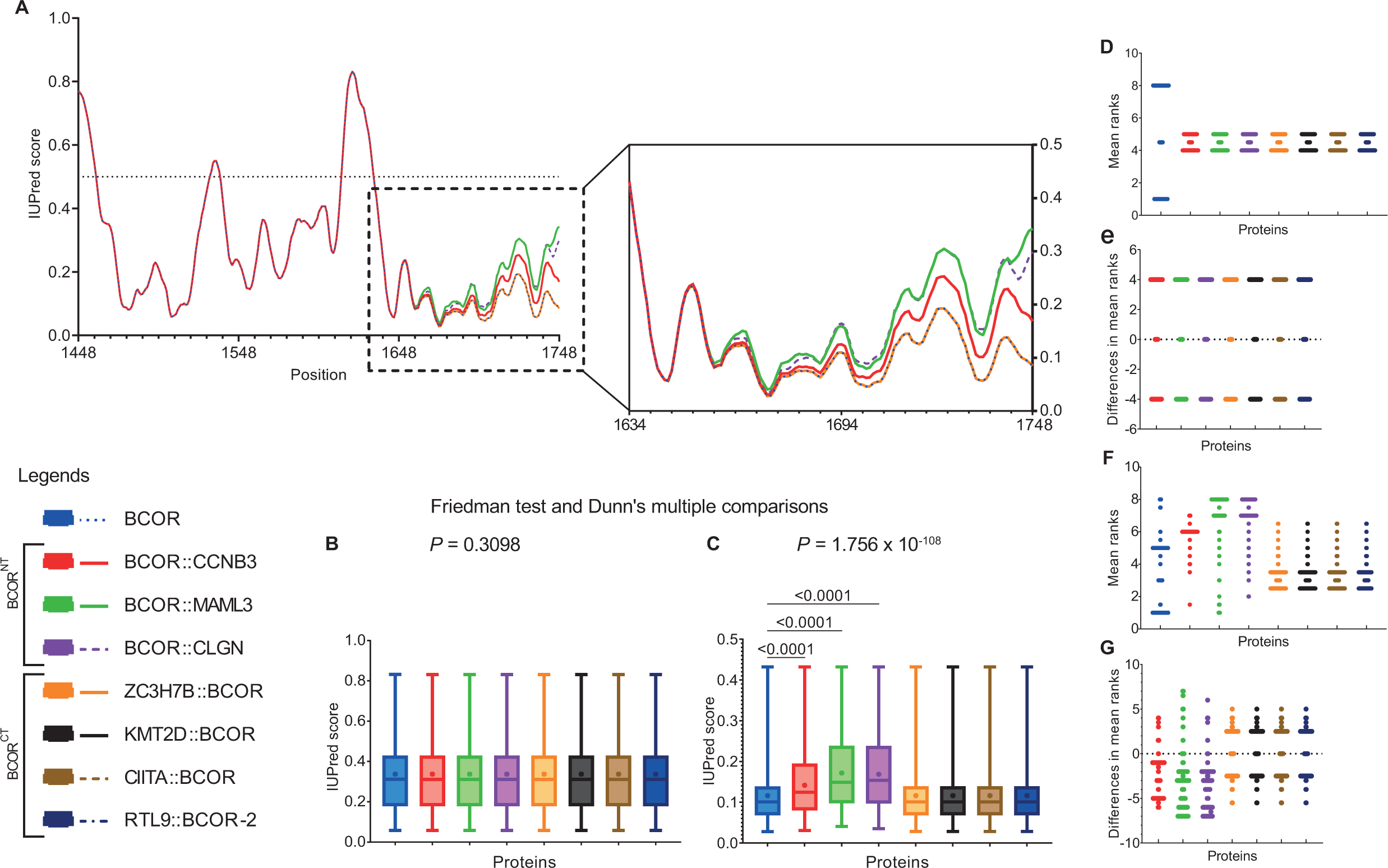
Comparison of the disorder propensities of wild-type BCOR and fusion proteins. The disorder propensities were calculated by using the IUPred3 online tool. The region^ANK+linker^ (encompassing 1448-1633 residues) and region^PUFD^ (encompassing 1634-1748 residues) of BCOR were investigated in the context of the BCOR and the fusion proteins. **(A)** IUPred3 scores for IDRs of BCOR and its fusion proteins in the coordinate system. **(B,C)**, Friedman’s test, and Dunn’s multiple comparisons with *P* values for BCOR and the fusion proteins (*n* = 8). The horizontal line shows the median, plus symbol (+) the mean value, and error bars represent the min-max. **(D,F)** Mean rank plot of Friedman’s test. (**E,G),** Friedman’s test’s differences in mean ranks between pairs of groups (BCOR vs. each fusion). Region^ANK+linker^ and region^PUFD^ are represented by (**A,B,D,E**) and (**A,C,F,G**) respectively. **BCOR^NT^**: BCOR is located at the N-terminal end of the fusion protein. **BCOR^CT^**: BCOR is located at the C-terminal end of the fusion protein.

Region^ANK+linker^ of wBCOR (containing 1448-1633 residues) consists of ANK repeats and a linker region, region^PUFD^ includes the PUFD domain (consisting of the 1634-1748 residues).

We did not find significant differences between the IUPred3 IDR scores of the proteins within the region^ANK+linker^ in at least two groups (Q = 8.265, *P* = 0.3098). However, significant differences were found within the region^PUFD^ in at least two groups (region^PUFD^: Q = 521.7, *P* < 0.0001). Dunn’s multiple comparison tests revealed significant rank means differences between the IUPred3 IDR scores of the BCOR and BCOR^NT^ proteins in both region^ANK+linker^ and region^PUFD^. There were significant differences in the rank means of BCOR vs. BCOR::CCNB3 (Z = 8.130, *P* < 0.0001), BCOR vs. BCOR::MAML3 (Z = 11.10, *P* < 0.0001), and BCOR vs. BCOR::CLGN (Z = 11.99, *P* < 0.0001). There was no statistically significant difference in the rank means between BCOR and ZC3H7B::BCOR, KMT2D::BCOR, CIITA::BCOR, and RTL9::BCOR-2 fusion proteins (**Fig. 3C ,D**).

The Z-score in Dunn’s multiple comparisons serves as a statistic that indicates the magnitude of the difference between the two groups being compared. In our analysis, the largest difference within the BCOR^NT^ group compared to the BCOR group was observed in the BCOR::CLGN fusion (Z = 11.99), followed by the BCOR::MAML3 fusion (Z = 11.10), and then the BCOR::CCNB3 fusion protein (Z = 8.13). Despite these substantial differences, it is noteworthy that none of its IDR scores reached the 0.5 disordered cut-off threshold. Consequently, despite notable differences in IDR scores in the region^PUFD^, the absence of IDR scores exceeding the disordered cut-off suggested that these regions retained a structured conformation.

According to the IPred3 prediction, the BCOR^NT^ fusions increased the disorder propensity in the region^PUFD^, potentially affecting the protein’s interaction with the PCGF1 RAWUL and KDM2B leucine-rich repeat (LRR) region. Such changes were not predicted for the members of the BCOR^CT^ group.

### Intermolecular Contacts, Binding Energy, and Structural Comparison

The PUFD domain remains fully intact after the fusion events, except in the case of BCOR::MAML3 where four C-terminal residues of this domain are lost (**Fig. 1**). Despite retaining the integrity of its sequence, the fusion events may potentially have an impact on the structure of the PUFD domain, which might alter its interactions with the RAWUL domain of PCGF1. To investigate these effects, we generated the structures of the complexes containing the RAWUL domain of PCGF1 and the PUFD domain of each fusion protein (excepting KMT2D::BCOR), by using AlphaFold2-multimer (AF2) (**Fig. 5B**). The structural analyses were limited to those regions that were present in the crystal structure of the complex of RAWUL and PUFD domains (PDB ID: 4hpl). The 4hpl crystal structure was the only available structural coordinate that has been determined by X-ray crystallography as well as made the investigation of protein-protein interactions possible because this structural file contains the RAWUL-PUFD dimer. We analysed the ranked 1 complexes using PRODIGY software to determine intermolecular contacts (ICs), ΔG and ΔΔG values in the RAWUL-PUFD complexes of the fusion proteins (**Fig. 4**). We compared the results to those of the 4hpl complex and only considered changes that exceeded the threshold (ICs: number in IC changes 2, NIS charged: 0.16%, NIS apolar: 0.25%; ΔΔG: 0.5).

**Figure 4.**
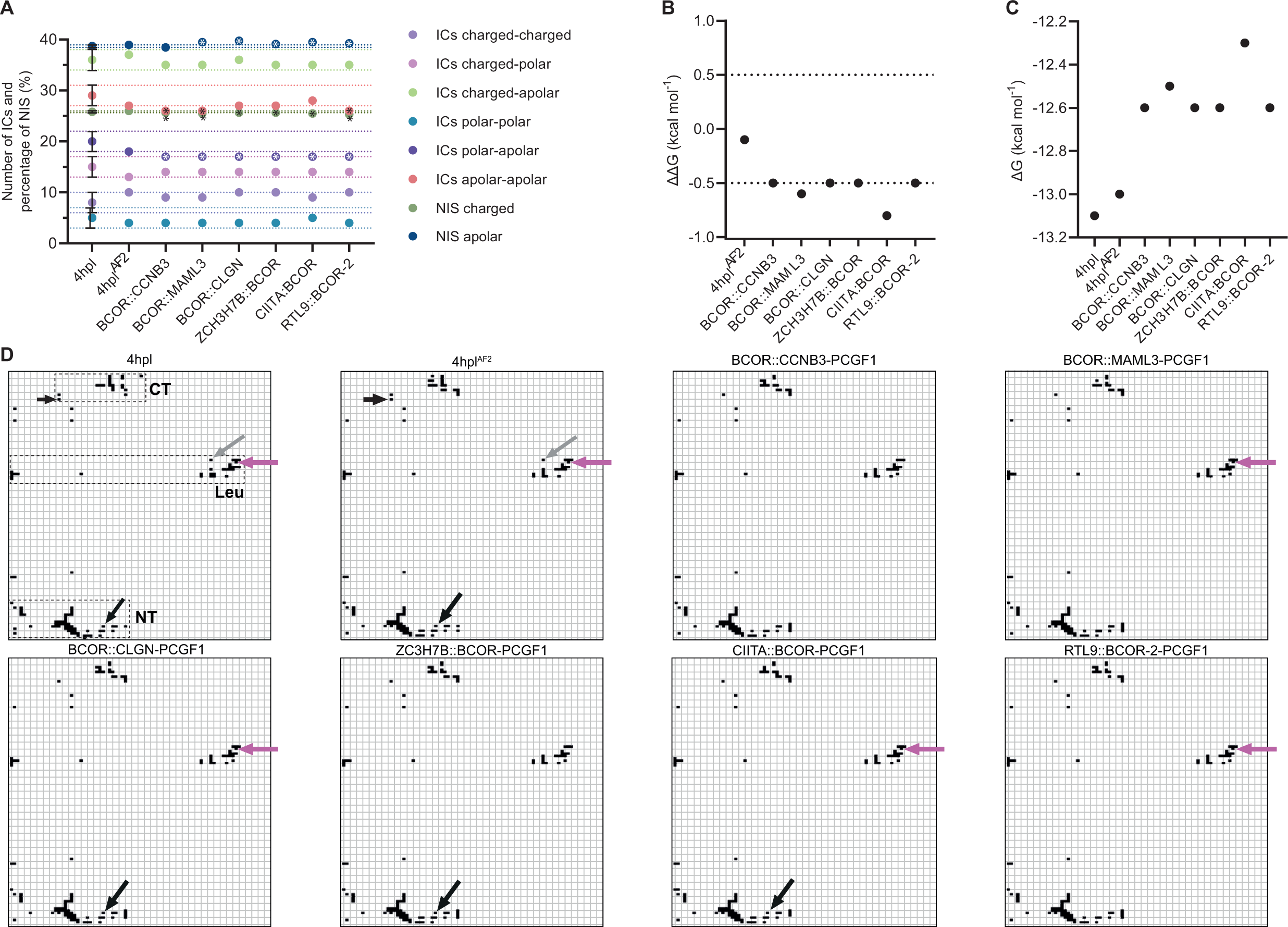
Intermolecular contacts (ICs) and binding affinity of RAWUL-PUFD dimers. **(A)** Intermolecular contacts were determined for the complexes containing the PUFD domain of the wild-type BCOR (4hpl and 4hpl^AF2^) or that of the fusion proteins. The prediction error range is indicated by the dotted line (+/-) and is derived from the difference between 4hpl and 4hpl^AF2^. **(B)** The binding affinity change of the predicted RAWUL-PUFD dimers in Gibbs free energy change (ΔΔG=ΔG_wild-type_-ΔG_fusion-type_). ΔΔG calculations were also performed for the 4hpl^AF2^. The dotted lines represent the 0.5 cut-off. **(C)** Binding affinity ΔG (kcal mol^-1^) of the protein complexes. The dotted lines indicate the threshold and the Asterisk (*) symbol the value out of the threshold in (a). **(D)** Contact maps were determined based on the RAWUL-PUFD complexes. Long black arrows show the IC between Phe1641 and Leu203 residues, this contact is absent if BCOR::CCNB3, BCOR::MAML3, and RTL9::BCOR-2 fusion proteins are in the complexes. Dotted circles indicate the NT, CT, and Leu interaction surfaces on the 4hpl contact map. The same interfaces are represented for all the complexes. Magenta arrows show the Lys1711-Pro245 IC which is not present in the BCOR::CCNB3 and ZC3H7B::BCOR complexes. Grey arrows show the Asp1712-Trp237 IC in the 4hpl and 4hpl^AF2^, this contact is missing in the case of each fusion protein complex. Short black arrows show the Gly1738-Asn189 IC which is absent from the complexes formed by the fusion proteins. The values obtained for the residues of RAWUL (167-254) and PUFD domains (1636-1748) are shown in the X and Y axes, respectively. **NIS**: percentage of the charged and apolar/polar non-interacting surface of the complex.

Ten different interaction types were analysed. It was discovered that four of them - charged-charged, polar-polar, charged-polar, and charged-apolar ICs - remained within the thresholds in the fusion protein group as compared to the wild-type (4hpl). The fusion complexes exceeded the threshold for the binding affinities changes, ICs polar-apolar, and percentage of the charged non-interacting surface (NIS charged). Only the BCOR::CCNB3, BCOR::MAML3, and RTL9::BCOR-2 fusions exceeded the threshold in the case of the apolar-apolar ICs. Additionally, the percentage of apolar NIS (NIS apolar) values remained unchanged except for BCOR::CCNB3 (**Fig. 4A**).

Our analysis showed that the predicted ΔGs decreased by at least 0.5 ΔΔG in all cases, with a mean ΔG of -12.533 kcal mol^-1^. The predicted ΔG values for the fusion proteins were as follows: CIITA::BCOR had the weakest binding affinity with a value of -12.30 kcal mol^-1^, while BCOR::MAML3 had a slightly stronger binding affinity with a value of -12.50 kcal mol^-1^. The complexes containing the PUFD domains of BCOR::CCNB3, BCOR::CLGN, ZC3H7B::BCOR, and RTL9::BCOR-2 fusion proteins exhibited highly similar ΔG values (approximately -12.6 kcal mol^-1^) (**Fig. 4C**) (**Table 5**).

**Table 5.** Binding affinities predicted for RAWUL-PUFD complexes. The column titles display the various PUFD regions or residues utilized to predict the binding affinity in the RAWUL-PUFD complex. The Complete PUFD structure shows the predicted ΔG values of the entire investigated RAWUL-PUFD dimer (RAWUL 167-177 and 185-254, as well as PUFD 1636-1748). The PUFD’s Interaction surfaces include the Interaction-Surface^NT/CT^ and Interaction-Surface^Leu^. The Interaction-surface^NT/CT^ consists of the residues in the regions 1636-1651 and 1738-1748, while the Interaction-surface^Leu^ contains the residues in the region 1704-1712. The ΔΔG (ΔΔG=ΔG_wild-type_-ΔG_fusion-type_) cut-off value was 0.5, to interpret the change in the binding affinity between 4hpl and the fusion complexes. Values that reach the cut-off are shown in bold. The values are in kcal mol^-1^. The lower ΔG value the higher the binding affinity. The stronger binding affinity may be due to the high ratio of binding residues to total residues of the "interaction surfaces" compared to the "complete PUFD structure".

|  | Complete<br>PUFD<br>structure | Interaction<br>surfaces | Interaction-Surface<sup>NT/CT</sup> | Interaction-Surface<sup>Leu</sup> | Leu1706 | Phe1639,<br>Phe1641 |
| --- | --- | --- | --- | --- | --- | --- |
| 4hpl | -13.1 | -13.8 | -11.2 | -8.3 | -5.0 | -6.3 |
| 4hpl<sup>AF2</sup> | -13.0 | -13.7 | -11.2 | -8.2 | -5.0 | -6.3 |
| BCOR::CCNB3-PCGF1 | **-12.6** | **-13.3** | -10.8 | -8.1 | -5.1 | -6.2 |
| BCOR::MAML3-PCGF1 | **-12.5** | **-13.0** | **-10.5** | **-7.8** | -5.0 | -5.9 |
| BCOR::CLGN-PCGF1 | **-12.6** | **-13.2** | **-10.7** | **-7.8** | -5.0 | -5.9 |
| ZCH3H7B::BCOR-PCGF1 | **-12.6** | **-13.3** | -11.0 | -7.9 | -5.1 | -6.3 |
| CIITA:BCOR-PCGF1 | **-12.3** | **-12.9** | **-10.5** | -8.0 | -5.0 | -6.1 |
| RTL9::BCOR-2-PCGF1 | **-12.6** | **-13.2** | **-10.7** | -8.0 | -4.9 | -6.1 |

To identify potential hotspots with altered interactions and binding affinity, we used MAPIYA to build contact maps. The analysis showed that the interactions between the RAWUL and PUFD domains were most dense in two regions (**Fig. 4C**). The first region of PUFD interacts with the previously described RAWUL β-sheet and loop binding surface (Junco *et al*, 2013) (**Fig. 5D**). The regions of PUFD that interacted with the binding surfaces of RAWUL were further investigated. These sites included the Val1636 residue, the N-terminal β-sheet (Phe1637-Ser1642) and its post-region (Glu1643-Asn1651), and the loop region (Gly1738, Ser1739) before the C-terminal β-sheet (Ser1740-Leu1744) and its post region His1745-Asp1748 (hereafter Interaction-Surface^NT/CT^) (**Fig. 5D**). The interaction surface that displayed a high frequency of ICs in the RAWUL-PUFD interaction was identified as the surface was containing the Leu1706 residue within the Ser1704-Asp1712 region (**Fig. 5D**). This area was referred to as Interaction-Surface^Leu^. Both interaction regions contain relevant ICs that we examined individually. The Interaction-Surface^NT/CT^ involves the Phe1639 and Phe1641 residues, which form hydrophobic contact with the RAWUL β-sheet binding surface residue Val206 (**Fig. 5F**). On the other hand, the Interaction-Surface^Leu^ includes the Leu1706 residue, which establishes contact with the RAWUL domain’s Leu Cage (Junco *et al*, 2013) (**Fig. 5E**). To highlight the interaction surfaces that predominantly contribute to the change in binding affinity, a comprehensive analysis was performed, examining them individually for changes in IC and changes in ΔG. This approach helps identify interaction surfaces that were crucial to the observed shifts in binding affinity, providing insights into the structural dynamics of the protein complexes.

**Figure 5.**
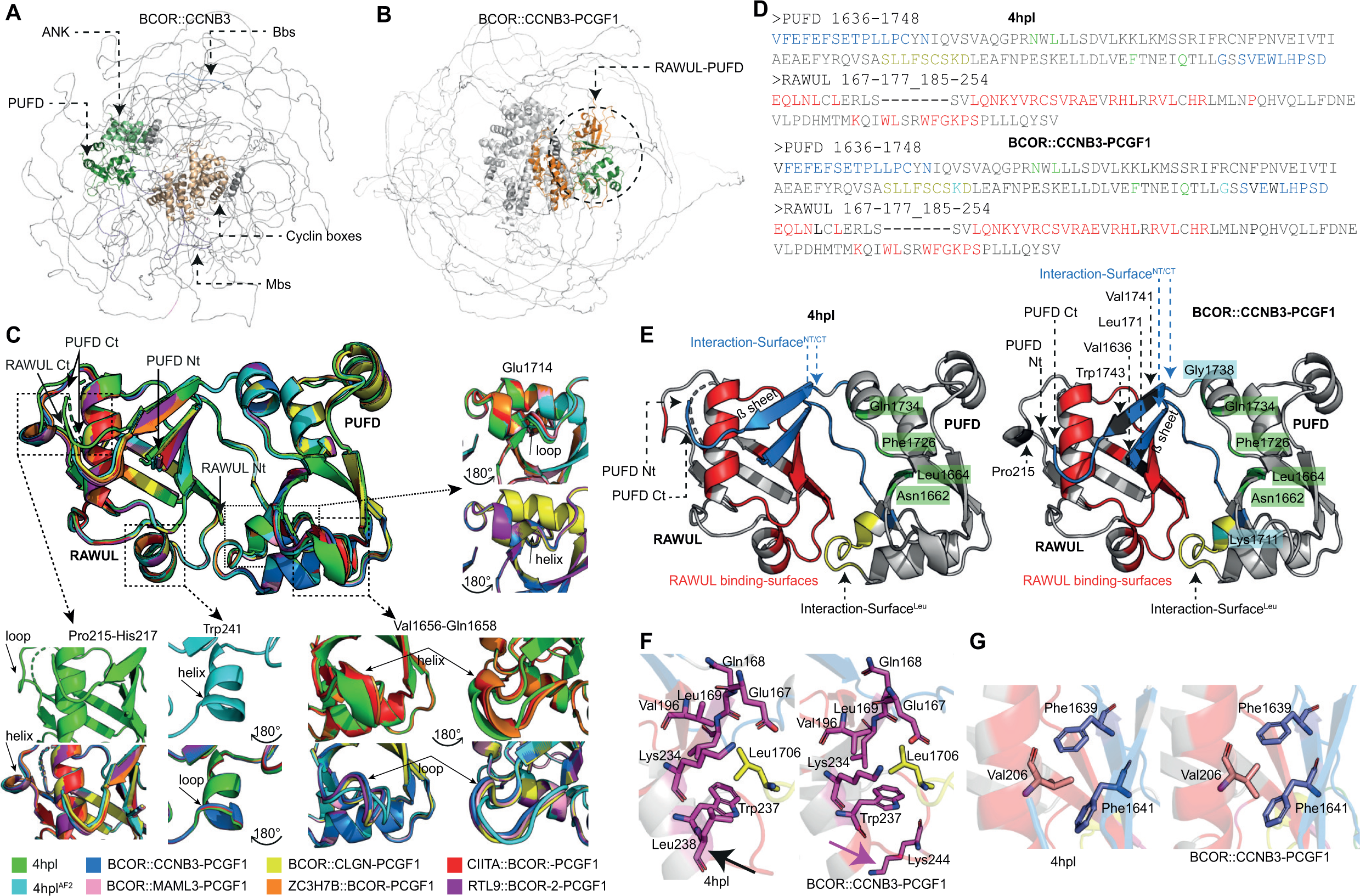
Structural analysis of fusion proteins and their complex with PCGF1. **(A)** AlphaFold2-predicted structure of BCOR::CCNB3. BCL6 binding site (Bbs, light blue), MLLT3 binding site (Mbs, purple), ANK rpt (rpt: repeats) (light green), PUFD domain (forest green), and Cyclin boxes (gold). The Mbs and Bbs are disordered regions and therefore difficult to recognize. **(B)** Structure of BCOR::CCNB3-PCGF1 complex. The BCOR::CCNB3 is shown with gray color, the PUFD domain is green, while the RAWUL domain of PCGF1 is orange and circled with a dashed line. **(C)** Aligned structures of the RAWUL-PUFD dimers. Structural differences were found at four sites among the dimers, specifically in the PUFD domain at Val1656-Gln1656 and Glu1714, and in the RAWUL domain at Pro215-His217 and Trp241. The structural variations across different dimers were compared by color-coding each, according to the legends. **(D,E),** 4hpl and BCOR::CCNB3-PCGF1 dimers and their amino acid sequences. Blue color marks the Interaction-Surface^NT/CT^ of PUFD, including the N-terminal (Nt) β-sheet (Phe1637-Ser1642) and its post-region (Glu1643-Asn1651), and the loop region (Gly1738, Ser1739) before the C-terminal (Ct) β-sheet (Ser1740-Leu1744) and its post region His1745-Asp1748. Red color marks the binding sites of RAWUL. Yellow color marks the Interaction-Surface^Leu^ of PUFD (yellow, 1704-1712). Green color marks residues in PUFD that are not part of the interaction surfaces but interact with RAWUL. The cyan color shows those PUFD residues of BCOR::CCNB3-PCGF1 that do not interact with the RAWUL domain. Black color marks BCOR::CCNB3-PCGF1 residues that are missing from the predicted domains (including 4hpl^AF2^). The color scheme of the amino acid sequences and the figures are consistent. **(F)** The PUFD Leu1706 (yellow stick) is surrounded by RAWUL residues (pink sticks) forming the Leu cage. In the BCOR::CCNB3-PCGF1’s Leu cage, the IC partner of PUFD Leu1706 residue changes from RAWUL Leu238 (black arrow) to Lys244 (magenta arrow). **(G)** PUFD Interaction-Surface^NT/CT^’s β-sheets (blue) Phe1639 and Phe1641 residues (blues sticks) interaction with RAWUL Val206 (white sticks).

The Interaction-Surface^NT^ interacts with 22 residues of the 167-210 region of RAWUL. This interaction surface of the RAWUL domain corresponds to the previously described β-sheet-and loop-binding surfaces (Junco *et al*, 2013). The PUFD domain of each fusion protein formed two new contacts with the RAWUL domain within the Interaction-Surface^NT^ which interactions were not present in the 4hpl and 4hpl^AF2^: Glu1640-Arg210 (salt bridge, hydrogen bond), and Ser1642-Arg210 (ion-dipole, hydrogen bond). Furthermore, contacts between the Asn1651 and Gln168 residues (dipole-dipole, hydrogen bond) contacts were also formed in the CIITA::BCOR complex. The Phe1641-Leu203 (hydrophobic) contact was not observed for the BCOR::CCNB3, BCOR::MAML3, and RTL9::BCOR-2 complexes, although, it was present in the complex formed between the wild-type proteins (4hpl). It is important to note that Val1636-Val197 (van der Waals force), Glu1640-Gln168 (ion-dipole, hydrogen bond), and Phe1641-Gln188 (π-π stacking) Leu1646-Leu171 (hydrophobic) contacts were absent from the predicted structures but were present in the 4hpl. In contrast, the Phe1639-Glu199 (anion-π stacking) interaction was found in the CIITA::BCOR-PCGF1 dimer, and the Phe1639-Arg210 (cation-π stacking) interaction was detected in the BCOR::CCNB3-, BCOR::MAML3-, BCOR::CLGN-PCGF1, RTL9::BCOR-2-PCGF1 complexes and both were found in the 4hpl^AF2^ and ZC3H7B::BCOR-PCGF1 structures but not in the 4hpl. The fact that Val1636-Val197, Glu1640-Gln168, Phe1641-Gln188, Leu1646-Leu171 were detected only in the 4hpl crystal structure, and Phe1639-Glu199, Phe1639-Arg210 were detected only in the 4hpl^AF2^ and in the dimers of the fusion proteins - to varying degrees - may be due to the rigid nature of the 4hpl structure and therefore these possible interactions were not identified. Prediction errors cannot be excluded either.

The Interaction-Surface^CT^ interacts with 9 residues of the 189-210 region of RAWUL. All fusion complexes gained a Glu1742-Leu207 (van der Waals force) interaction and an His1745-His202 (ionic repulsion) IC except BCOR::MAML3. However, they all lost the Gly1738-Asn189 (van der Waals force) and Pro1746-Arg205 (van der Waals force) contacts. Furthermore, the contact between His1745 and Arg205 (ionic repulsion) was absent only from the BCOR::CCNB3 and CIITA::BCOR fusion proteins.

Similar phenomena to those observed in the Interaction-Surface^NT^ were also observed in the 4hpl^AF2^ structure, where some residue-residue interactions were missing. Additionally, the predicted structures revealed new ICs that may not have been previously captured in the 4hpl dimer or may have been caused by the AF2 method. The Trp1743-Val206 (hydrophobic), Trp1743-His209 (cation-π stacking, π-π stacking, hydrogen bond), Leu1744-His209 (van der Waals force), Ser1747-His209 (electrostatic ion-dipole, hydrogen bond, dipole-π stacking) Asp1748-His209 (salt bridge, hydrogen bond, anion-π stacking), Asp1748-Pro215 (van der Waals force) were absent from the predicted structures and were only found in the 4hpl. However, the Val1741-Arg210 (van der Waals force), Glu1742-His209 (salt bridge, hydrogen bond, anion-π stacking), Pro1746-His202 (van der Waals force) interactions were found in the BCOR::MAML3-, CIITA::BCOR-PCGF1 dimers, additionally also the Pro1746-Arg201 (van der Waals force) contact in the BCOR::CCNB3-, BCOR::CLGN-, ZC3H7B::BCOR-PCGF1, RTL9::BCOR-2-PCFG1 complexes, and even the Glu1742-Arg205 (salt bridge, hydrogen bond) and the Asp1748-His202 (salt bridge, hydrogen bond, anion-π stacking) ICs were discovered in the 4hpl^AF2^ structure, but none of these were in the 4hpl.

The Interaction-Surface^Leu^ interacts with 13 residues of the 167-246 region of the RAWUL domain. None of the predicted structures showed the Leu1706-Leu238 (hydrophobic) contact unlike the wild-type PUFD-RAWUL complex (4hpl). The fusion complexes gained a hydrophobic IC replacement, resulting in the Leu1706-Lys244 contact. Furthermore, in the fusion complexes, CCNB3::BCOR lost contact between Lys1711 and Pro245 (van der Waals force), ZC3H7B::BCOR also lacked contact between Lys1711-Pro245 and Phe1707-Glu167 (anion-π stacking), and CIITA::BCOR no longer had contact Asp1712-Trp237 (anion-π stacking, π-π stacking, and hydrogen bonding) (**Fig. 4C**). The same problem as before was encountered in this interaction surface. However, only two ICs, Leu1706-Leu238 (hydrophobic) and Cys1709-Ser246 (electrostatic dipole-dipole, hydrogen bond) were missing from the predicted structures and were found in the 4hpl. However, all AF2 structures contained the Phe1707-Trp237 (π-π stacking, hydrophobic) interactions while the 4hpl did not.

To compare the changes in ΔG of the key regions between the wild-type and the fusion protein complexes, we predicted the binding energy across the interaction surfaces, including the Interaction-Surface^NT/CT^ and the Interaction-Surface^Leu^ (**Fig. 5D**, **Table 5**). The binding affinity ΔG and values calculated for the interaction surfaces in the case of 4hpl and 4hpl^AF2^ structures were almost identical and were -13.8 and -13.7 kcal mol^-1^, respectively. The fusion complexes showed considerable differences, corresponding to the 0.5 ΔΔG cut-off value as compared to the wild-type structures. The complex of PCGF1 with BCOR::CCNB3 or ZC3H7B::BCOR had the strongest ΔG of -13.3 kcal mol^-1^, while the BCOR::CLGN-PCGF1 and RTL9::BCOR-2-PCGF1 dimers had a ΔG of -13.2 kcal mol^-1^. The BCOR::MAML3-PCGF1 complex (-13.0 kcal mol^-1^) and the CIITA::BCOR-PCGF1 complex (-12.9 kcal mol^-1^) exhibited the weakest binding affinity values.

Predictions were made for Interaction-Surface^NT/CT^, as well. The wild-type 4hpl structure exhibited a ΔG of -11.2 kcal mol^-1^, while all fusion protein-containing complexes showed a lower binding affinity, indicating weakened interaction at these sites between the RAWUL and PUFD domains. Only the BCOR::CCNB3-and ZC3H7B::BCOR-PCGF1 alterations were below the cut-off, these fusion proteins showed the most moderate decrease of binding affinity (3.6% and 1.8%, respectively). In contrast to this, ≥0.5 ΔΔG was predicted for the other complexes, and the most notable decrease (6.2%) was predicted for BCOR::MAML3-and CIITA::BCOR-PCGF1. Interestingly, the binding affinity of the RAWUL-PUFD dimer was lower in the case of each fusion protein as compared to the wild-type complex, although, the fusion complexes exhibited a higher number of ICs (**Table 5**).

The Phe1639-Val206 and Phe1641-Val206 interactions (hydrophobic) between the Interaction-Surface^NT^ of PUFD and the ß-sheet-binding surface of the RAWUL domain (Junco *et al*, 2013) were present in all fusion complexes. The results showed that the ΔG is decreased upon protein fusion, except for ZC3H7B::BCOR-PCGF1. However, none of the fusion complexes’ ΔΔG met the cut-off value (**Fig. 5D**). The largest decrease in binding affinity (6.35%) was calculated for the BCOR::MAML3-and BCOR::CLGN-PCGF1 dimers (**Table 5**).

Binding affinity prediction was performed for the Interaction-Surface^Leu^, as well (**Fig. 5C**). We observed that the interaction between the fusion proteins and the RAWUL domain was notably weaker at this site in two cases. As compared to the -8.3 and -8.2 kcal mol^-1^ ΔG calculated for 4hpl and 4hpl^AF2^, respectively, the fusion proteins showed weakened binding affinity. Two ΔΔG values exceeded the cut-off, the BCOR::MAML3-and BCOR::CLGN-PCGF1 complexes (0.5 ΔΔG), with a 6.41% change in ΔG (**Table 5**).

The binding affinity between the PUFD Leu1706 and the Leu cage (Junco *et al*, 2013) of the RAWUL domain was also predicted. The values that were predicted for 4hpl and 4hpl^AF2^ dimers as well as for the fusion complexes were almost identical, indicating that the hydrophobic IC swap probably did not affect the binding affinity of Leu1706 to its “cage” (**Table 5**).

The predicted structures either lacked or contained interactions that were absent or present in 4hpl. This may be because the X-ray crystallography captures only a snapshot or an averaged representation of a dynamic protein (Srivastava *et al*, 2018), as a result, the 4hpl^AF2^’s 3D state may also be a valid structure despite the different IC landscape. This was supported by the almost identical ΔG values between the two wild-type structures, 4hpl and 4hpl^AF2^, which suggest that the binding affinity was not affected. While in the fusion dimers, where additional IC gains but mostly losses were detected compared to the 4hpl^AF2^, the ΔG values were notably larger.

To further compare the protein structures, we aligned them with 4hpl as the reference (**Fig. 5C**). We observed minor changes in the secondary structures of the proteins. Specifically, the PUFD domain at residues Val1656-Gln1658 exhibits a helical conformation in the 4hpl structure, which is also observed in the CIITA::BCOR-and ZC3H7B::BCOR-PCGF1 complexes. However, in 4hplAF2, BCOR::CCNB3-, BCOR::MAML3-, BCOR::CLGN-PCGF1, and RTL9::BCOR-2-PCGF1, these residues adopt a loop conformation. It is important to note that the variation includes 4hpl^AF2^, which is not involved in fusion events, and this region does not interact with the RAWUL domain. Therefore, based on the binding affinity results, this change may have no significant consequences and could represent an alternative conformation not captured in 4hpl. Another minor alteration was observed in the PUFD domain at the Glu1714 residue, following the Ser1710-Leu1713 helical section. In the BCOR::CCNB3-, BCOR::MAML3-PCGF1, and RTL9::BCOR-2-PCGF1 complexes, this residue adopts a helical secondary structure, extending the Ser1710-Leu1713 helix. However, in the 4hpl, 4hpl^AF2^, BCOR::CLGN-, CIITA::BCOR-, and ZC3H7B::BCOR-PCGF1 dimers, it assumes a loop conformation. Despite its proximity to the Interaction-Surface^Leu^, there is no direct link between changes in intermolecular contacts and alterations in secondary structure.

Our analysis revealed two alterations in the RAWUL domain of the protein complexes. Firstly, in the 4hpl structure, the Pro215-His217 residues adopt a loop conformation, whereas in predicted structures, including 4hpl^AF2^, a helical secondary structure is observed. The predicted PUFD-RAWUL dimers may have changed due to the absence of the Ser1748-Pro215 intermolecular contact. This change could have potentially affected the position of the PUFD C-terminus (His1745-Asp1748) and influenced the secondary structure conformation near Pro215-His217. Furthermore, the 4hpl^AF2^ structure showed a specific alteration where the Trp241 residue changed conformation from a loop to a helix. Despite interacting with PUFD’s Ser1708 residue, this change may not significantly affect the interaction between the two domains.

The fusion complexes showed a decrease in ΔΔG and minor structural differences between the interaction surfaces of PUFD and the RAWUL domain. We have identified the origin of the decrease in binding affinity to the Interaction-Surface^NT/CT^ and the Interaction-Surface^Leu^. Notable decreases were found in the BCOR::MAML3-PCGF1 and BCOR::CLGN-PCGF1 complexes in both regions, as also in the Interaction-Surface^NT/CT^ in the case of the CIITA::BCOR-PCGF1, RTL9::BCOR-2-PCGF1 complexes. The interactions between Leu1706 and RAWUL’s Leu cage, as well as the interaction between PUFD’s Phe1639 and Phe1641 with Val206, did not meet the cut-off for ΔΔG values. However, it is important to note that for the Phe1639-and Phe1641-Val206 interactions, although they did not meet the cut-off value, a -0.4 ΔΔG was observed in two fusion complexes. This means a 6.35% decrease in binding affinity, indicating a subtle but relatively large ΔΔG in the binding affinity between these residues.

### Statistical binding affinity comparison of the RAWUL-PUFD dimers

We conducted 3D structure predictions of the fusion proteins using AF2, along with one prediction of the wild-type BCOR, in complex with PCGF1. Subsequently, we analysed and emphasized the changes in ΔG and ICs within the complexes, comparing the fusion-type dimers to the wild-types. However, statistical comparison between these groups posed a challenge. To address this, we used five predicted structures for each AF2 prediction. It is important to note that each prediction resulted in five structures, but for BCOR::MAML3-PCGF1 one prediction did not include the RAWUL-PUFD dimer.

The complexes were used to predict the binding affinity between the dimers and to compare the fusion-type complexes to the wild types (4hpl, 4hpl^AF2^). The statistical comparison to identify significant differences between the ΔG results of the groups was conducted using Ordinary one-way ANOVA and as a post hoc test of Tukey’s multiple comparisons were applied.

First, we analysed the whole studied dimers (RAWUL: 167-177, 185-254; PUFD: 1636-1748) and found a statistically significant difference in the mean ΔG values between at least two groups (F(7, 27) = [7.448], *P* <0.0001). Tukey’s multiple comparison tests found that the mean ΔG values were significantly different between 4hpl vs. BCOR::CCNB3-PCGF1, BCOR::MAML3-PCGF1, BCOR::CLGN-PCGF1, CIITA-BCOR-PCGF1 and between 4hpl^AF2^ vs. BCOR::CCNB3-PCGF1, BCOR::MAML3-PCGF1, BCOR::CLGN-PCGF1, ZC3H7B::BCOR-PCGF1, CIITA-BCOR-PCGF1 (*p* values on **Fig. 6A**). There was no statistically significant difference in the mean ΔG between 4hpl vs. 4hpl^AF2^ (*P* = 0.9910), 4hpl vs. ZC3H7B::BCOR-PCGF1 (*P* = 0.0964), 4hpl vs. RTL9-BCOR2-PCGF1 (*P* = 0,1646) and 4hpl^AF2^ vs. RTL9-BCOR2-PCGF1 (*P* = 0.0501). Also, there was no statistically significant difference in the mean ΔG values between the fusion-type complexes (**Fig. 6A**).

**Figure 6.**
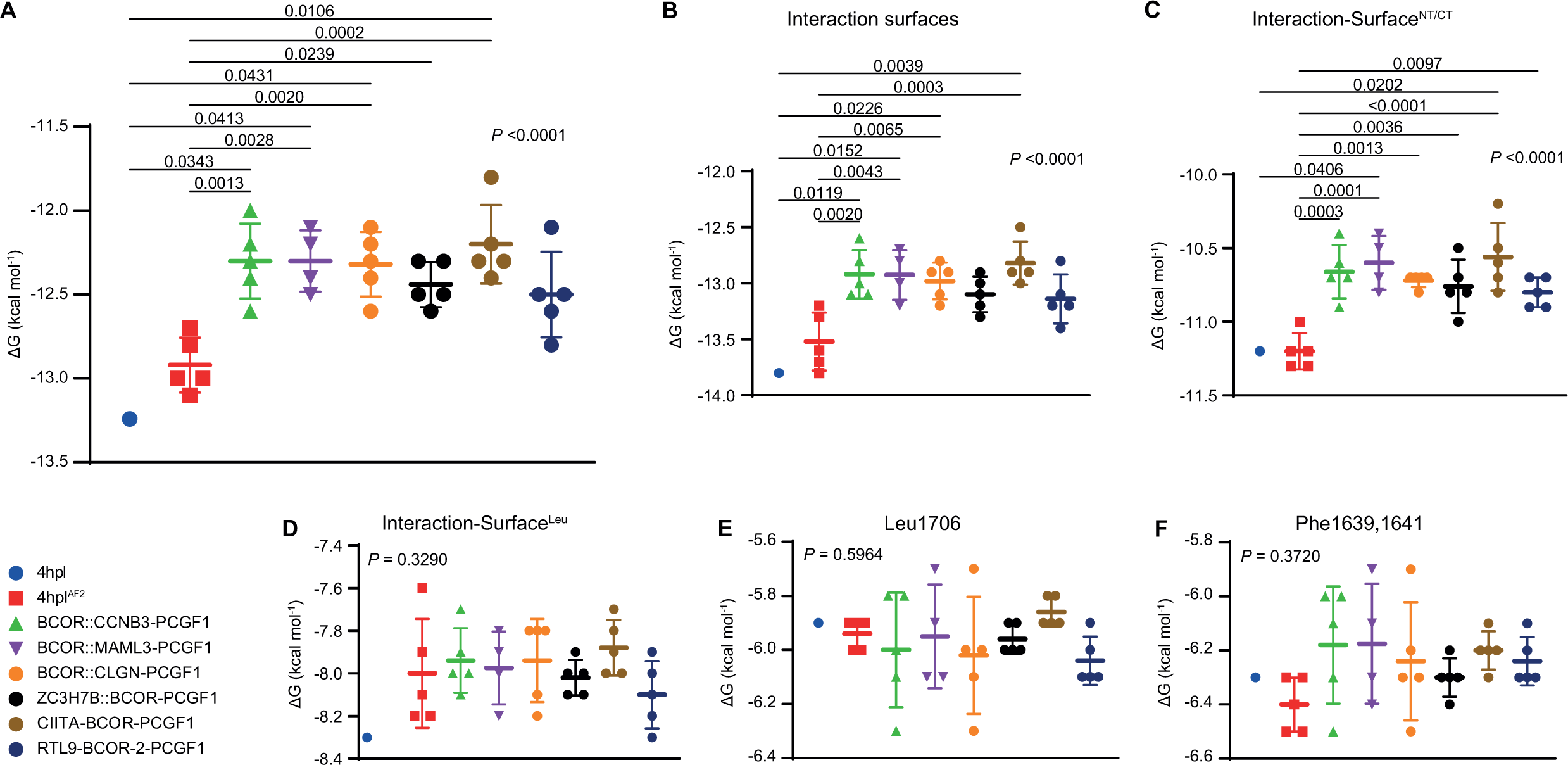
Analysis of binding affinity variations across PUFD interaction surfaces in *BCOR*-Rearranged Sarcomas. Statistical comparison analysis (Ordinary One-way ANOVA and Tukey’s multiple comparison test) with *P* values was conducted on the dimer’s (*n* = 8) binding affinity values. Error bars denote the standard deviations and the mean values. The six graphs depicted in the figure represent distinct scopes of PUFD in the context of the RAWUL domain. **(A)** The entire studied RAWUL and PUFD domain (RAWUL: 167-177, 185-254; PUFD: 1636-1748) **(B)** Interaction surfaces in the dimer with RAWUL. **(C)** Interaction-Surface^NT/CT^ in dimer with RAWUL. **(D)** Interaction-Surface^Leu^ in the dimer with RAWUL. **(E)** Leu1706 in the dimer with RAWUL. **(F)** Phe1639, 1641 residues in dimer with RAWUL. Significant differences between the eight PUFD groups were observed in cases of **(A-C)**. Post hoc analyses revealed significant results, indicated by braces, with the *P* value considered significant at *P* < 0.05. The **Interaction surfaces** include Val1636-Asn1651, Ser1704-Asp1712, and Gly1738-Asp1748. **Interaction-Surface^NT/CT^** consists of Val1636-Asn1651 (N-terminus) and Gly1738-Asp1748 (C-terminus). **Interaction-Surface^Leu^** specifically refers to the region encompassing Ser1704-Asp1712. **Leu1706** represents only the Leu1706 residue in the dimer with RAWUL, while **Phe1639,1641** pertains solely to Phe1639 and Phe1641 residues in the dimer with RAWUL.

To pinpoint specific areas of change, we used ordinary one-way ANOVA tests on various scopes. Initially, we concentrated on the interaction surfaces of PUFD (Val1636-Asn1651, Ser1704-Asp1712, and Gly1738-Asp1748), then proceeded to analyse individual components, such as Interaction-Surface^NT/CT^ (Val1636-Asn1651, Gly1738-Asp1748) and Interaction-Surface^Leu^ (Ser1704-Asp1712), and finally examined previously identified key residues (Leu1706; Phe1639 and Phe1641) (Junco *et al*, 2013) in complex with the RAWUL domain.

There was a statistically significant difference in the mean ΔG values of the interaction surfaces (F(7, 27) = [7.095], *P* <0.0001) (**Fig. 6B**), as well as the Interaction-Surface^NT/CT^ (F(7, 27) = [8,588], *P* <0.0001) (**Fig. 6C**) between at least two groups. Tukey’s multiple comparison test results in **Fig. 6D, c**. and **Table EV1**.

We found no statistically significant difference in the mean ΔG values of the Interaction-Surface^Leu^ (*P* = 0.3290) (**Fig. 6D**), the Leu1760 (*P* = 0.5964) (**Fig. 6E**), and the Phe1639, 1641 (*P* = 0.3720) residues (**Fig. 6F**).

Statistical comparisons between wild-type and fusion-type proteins revealed a significant decrease in binding affinity in most cases. Notably, significant differences in mean ΔG values were observed for all fusion complexes compared to 4hpl, except for ZC3H7B::BCOR-PCGF1 and RTL9::BCOR-2-PCGF1. However, even in these cases, the mean differences (ΔΔG) were notable, at -0.66 and -0.6, respectively. Interestingly, 4hpl^AF2^ showed no significant difference compared to RTL9-BCOR-2-PCGF1, with a mean difference of -0.42 (**Fig. 6A**).

When narrowing down the analyses to the interaction surfaces of BCOR, fewer significant differences were observed. However, significant disparities were found between 4hpl^AF2^ and all fusion-type complexes in Interaction-Surface^NT/CT^. Conversely, no significant differences were observed in Interaction-Surface^Leu^, Leu1706, and Phe1639,1641 residues.

It is noteworthy that regardless of the scope, there were no significant differences between the 4hpl and the 4hpl^AF2^ complexes. This suggests that the observed differences in the fusion proteins were not merely the result of prediction error. A limitation of the analyses was that while 4hpl was a single experimentally determined structure, the predicted dimers had five versions (BCOR::MAML3-PCGF1 only had four), and there were prediction quality differences among them.

## Discussion

*BCOR* was first identified as a novel interacting corepressor of *BCL-6* that enhances BCL-6-mediated transcriptional repression (Huynh *et al*, 2000). Later, the specific BCL6 binding motif was found at the BCOR’s 498-514 site (Ghetu *et al*, 2008). Additionally, it was discovered that the BCOR directly interacts with the transcriptional regulator *AF9* (*MLLT3*) (Bushweller *et al*, 2018) and binds to chimeric MLL–AF9 proteins (MLL: mixed lineage leukemia), although, only some isoforms of BCOR bind AF9 (Srinivasan *et al*, 2003). BCOR is a known part of the PRC1.1. During the assembly of this complex, an interaction is formed between the BCOR’s PUFD domain and the PCGF1 protein’s RAWUL domain. Additionally, the PUFD’s ANK repeats and the linker region must also interact with the KDM2B’s C terminus (Blackledge *et al*, 2014; Wong *et al*, 2020).

A new subset of gene fusion in bone sarcoma was described first in 2013 by Pierron *et al* (Pierron *et al*, 2012). Since then six additional fusion genes have also been identified (Kao *et al*, 2018; Pierron *et al*, 2012; Specht *et al*, 2016; Yoshida *et al*, 2020; Vasella *et al*, 2022; Chang Chien *et al*, 2023) and classified into the third distinct subset of "undifferentiated small round cell sarcomas of bone and soft tissue" (Pierron *et al*, 2012; Sbaraglia *et al*, 2020). The pathological features have been thoroughly investigated, but the exact sequence assembly and protein-level functional analysis of the fusion proteins have not been performed yet. In this study, we utilized *in silico* tools to explore the properties of these uncommon fusion proteins. For this purpose, we aimed to analyse the sequences of the known *BCOR* rearrangements in sarcomas and compare the fusion proteins’ domain architectures (**Fig. 1**) and physicochemical characteristics (**Table 1**, **Fig. 2**), as well as to predict and analyse the PANTHER GO terms (**Table 3**), signal peptides, intramolecular localizations (**Table 4**), and IDRs (**Fig. 3**). The 3D structures of BCOR-PCGF1 complexes were also investigated, with a special emphasis on the ICs and binding energy between the RAWUL-PUFD domains (**Fig. 4**), of BCOR and partner proteins.

The seven fusion proteins were classified into two groups based on the localization of BCOR in the fusion protein: BCOR^NT^ and BCOR^CT^. The characteristics of the fusion proteins are described as follows.

The first member of the BCOR^NT^ group was the BCOR::CCNB3 which is a 3038 AA long protein. This fusion protein contains the whole sequence of BCOR and the CCNB3 protein truncated at its N terminus. The consequence of chromosomal rearrangement was the loss of 1-113 region of CCNB3 protein. This region contains the destruction box motif which may act as a recognition signal for degradation *via* the ubiquitin-proteasome pathway, and in addition, the mitotic destruction of cyclin B3 is also dependent on the destruction box sequence (Nguyen *et al*, 2002). The two cyclin boxes at the C terminus of CCNB3 are not lost upon the fusion.

The second BCOR^NT^ was the 2734 AA long BCOR::MAML3 fusion protein. In this protein, the BCOR lost only four C-terminal residues, so the truncation does not affect its PUFD domain. But on the other hand, the MAML3 lost its 1-157 region containing the N-terminal MamL-1 domain, which is responsible for the interaction with the ankyrin repeat region of the Notch proteins such as NOTCH1, NOTCH2, NOTCH3, and NOTCH4 (Chiang *et al*, 2006; Liu *et al*, 2009; Nam *et al*, 2006; Wu *et al*, 2007).

The third BCOR^NT^ protein was the BCOR::CLGN, in which the BCOR sequence remains intact while calmegin loses almost half of its sequence, including most of its calreticulin sequence. This loss is likely to have an impact on the function of calmegin and, consequently, on male fertility (Ikawa *et al*, 1997).

In the case of ZC3H7B::BCOR, belonging to the BCOR^CT^ group, the BCOR protein lost its Bbs site and, consequently, the role of the corepressor function with BCL6. Although ZC3H7B retained its TPR and LD motifs, it lost its C2H2-and C3H1-type zinc fingers. ZC3H7B recognizes the hairpins of miR-7-1, miR-16-2, and weaker miR-29a. Knockout of ZC3H7B in HEK293 cells reduced mature miR7 levels (Treiber *et al*, 2017). The truncation of the protein and loss of its C2H2 and C3H1 zinc fingers, which may interact with RNA (Lai *et al*, 1999; Carballo *et al*, 1998; Brown, 2005), could potentially impact its interaction with miR-7.

Due to the fusion of *KMT2D* and *BCOR* genes, the resulting chimera protein does not contain the Bbs of the BCOR. Consequently, the protein has lost its BCL6 corepressor function. Additionally, due to its C-terminal truncation, the second ePHD, FYRN/C, and SET domains of KMT2D are missing from the fusion protein. The lost ePHD may have had a DNA binding function (Lee *et al*, 2023) while the FYRN/C and SET domains are crucial for protein-protein interactions within the COMPASS complex. This complex is crucial for the methylation of histone H3 at lysine 4 (H3K4) and plays an essential role in gene regulation and development (Cho *et al*, 2007).

Regarding the CIITA::BCOR fusion, the BCOR lost its Bbs while the CIITA protein lost its main part (146-1130) excepting the N-terminal region; this change may potentially impact the protein’s self-association (Linhoff *et al*, 2001) and the transcriptional activation of the major histocompatibility complex class II (León Machado & Steimle, 2021).

RTL9 is also known as sushi-ichi retrotransposon homolog 10 (SIRH10) or retrotransposon Gag domain-containing protein 1 (RGAG1). This protein belongs to the family of the so-called gag-like proteins whose domains are homologous to those of retroviral and retrotransposon polyproteins including the capsid structural protein (Ishino *et al*, 2023; Campillos *et al*, 2006). The 1169-1316 region of RTL9, close to its C terminus, encompasses both the N-and C-terminal subdomains (NTD and CTD) of the capsid-like domain. This domain shares a high structural similarity with the capsid domain of the human immunodeficiency virus and with the capsid-like domain of other mammalian gag-like proteins, such as activity-regulated cytoskeleton-associated proteins (Arc/Arg3.1). These proteins and other gag-like proteins including the members of the RTL family such as RTL1 (also known as PEG11) and RTL2 (also known as PEG10 or RGAG3) are known to have the ability to self-assemble into viral capsid-like particles, via oligomerization of the capsid-like domains (Segel *et al*, 2021). The intact RTL9 protein may also be potentially able for self-assembly, but this has not been proved experimentally so far. The RTL9::BCOR-2 fusion protein contains only the 1-1199 residues of the RTL9, almost the entire capsid-like domain of RTL9 is deleted upon fusion. Consequently, the RTL9 cannot facilitate the oligomerization of the fusion protein via its capsid-like domain.

The physicochemical properties of the fusion proteins were investigated and compared to those of the wild-type proteins (**Table 2**). The study results indicate that the mean change across all cases was 30.90%, while the |mean| was 46.64%. The BCOR^NT^ and BCOR^CT^ groups were examined separately, revealing that the mean change for the BCOR^NT^ group was 58.03%, with an absolute mean of 55.25%. The BCOR^CT^ group had a mean change of 10.54% and an absolute mean change of 40.18%. The average GRAVY values of fusion proteins decreased by 35.95% compared to wild-type proteins, except for CIITA and RTL9 proteins, which showed a decrease of 305.10% and 198.54%, respectively. Excluding these two proteins, the average GRAVY value increased by only 0.03%. Compared to the wild-type BCOR protein, the fusion proteins showed an increase in GRAVY values of 2.75%. The highest increase was observed in BCOR^CT^ alone, with a value of 5.11%. However, the GRAVY values did not reach the positive range, indicating that the protein structures could still be predicted to be globular (Kyte & Doolittle, 1982). The AI values remained largely unchanged. The II of the proteins increased by 7.4%. In the case of BCOR, the II showed an average increase of 10.71%. Additionally, for BCOR^NT^, the increase was 4.35%, while for BCOR^CT^, it was 15.49%. The increase in the II indicates a decrease in protein stability and a corresponding decrease in their *in vivo* half-life (Guruprasad *et al*, 1990). After the fusion event, the MAML3 protein showed the most substantial mean change of 124.69%, while RTL9 exhibited the most significant |mean| change of 131.18% when comparing the wild-type and fusion proteins. On the other hand, ZC3H7B had the smallest mean change of 0.47%, and KMT2D had the smallest |mean| change of 3.56%.

The Gravy, AI, -R(%), and +R(%) values showed a decrease while II increased for the fusion proteins. This led to an increase in protein hydrophobicity, a decrease in protein thermostability, a decrease in the relative value of charged amino acids, and an increase in the potential for protein degradation and denaturation (**Fig. 2**).

The predicted GO terms of the fusion proteins revealed that BCOR has a dominant role in the fusion proteins compared to other wild-type partners. A BCOR’s negative regulation of transcription by RNA polymerase II (GO:0000122), transcription corepressor activity (GO:0003714), and nucleus (GO:0005634) terms conserved in the fusion proteins expect of KMT2D::BCOR. This invariance is likely since BCOR has not lost any functional domains other than the lost Bbs domains of the BCOR^CT^ group, and its partners suffered major truncations (**Fig. 1**). KMT2D may have retained its terms because its N-terminal part at amino acid 4467 is intact. The retention of PANTHER GO terms in BCOR and KMT2D does not guarantee that the function of the proteins has not been affected by the fusion event. Limited information on RTL9 and so on the RTL9::BCOR-2 proteins may explain the lack of results (**Table 3**).

The proteins were screened for signal peptides using SignalP 6.0. Only calmegin was found to have a 99.92% probability of containing secreted signal peptides, while BCOR::CLGN did not. DeepLoc 2.0 was used to determine the intracellular localization of the proteins. The results indicate that almost all fusion proteins may be found in the nucleus, except BCOR::CLGN which may located in the endoplasmic reticulum. Analyses for GPI-anchoring signals were performed using the NetGPI-1.1 predictor, but no results were obtained for any of the proteins. The results overall were consistent with the UniProt and The Human Protein Atlas databases for wild-type proteins, suggesting that the predicted location of the fusion proteins was accurate (**Table 4**). These results are consistent with the PANTHER GO term cellular component, which also indicates nuclear localization for most of the fusion proteins. Only for the BCOR::CLGN, we observed ER localization using DeepLoc 2.0, but nuclear localization using PATHER GO terms. The CLGN is known to contain an N-terminal hydrophobic signal peptide (Watanabe *et al*, 1994) at predicted position 1-19 (Chang Chien *et al*, 2023), which is responsible for ER localization, and a hydrophilic C-terminus with a transmembrane domain and ER retention signal (Ikawa *et al*, 1997). As a result of the fusion event, the N-terminal hydrophobic signal is lost with the first 295 AA of the protein, but the C-terminal sites remain intact (1931–1952 residues, according to BCOR::CLGN). The N-terminal hydrophobic signal peptide is crucial for directing calmegin protein to the ER. Without this signal peptide, calmegin may not be correctly targeted to the ER, which can affect its proper localization and function within the cell. The C-terminus of CLGN functions to anchor the protein to the ER membrane and ensure its retention within the ER for its chaperone functions. This signal is crucial for maintaining calmegin’s localization within the ER, where it plays an important role in protein folding, quality control, and interactions related to spermatogenesis and sperm-egg interactions(Ikawa *et al*, 1997, 2001). DeepLoc 2.0 identified the C-terminus of CLGN and located it in the ER, despite the absence of the N-terminus. The KMT2D and KMT2D::BCOR’s cellular component term is MLL3/4 complex (GO:0044666) which is a histone methyltransferase complex located in the nucleoplasm of the nucleus. No literature data were available for RTL9::BCOR-2.

*BCOR*-rearranged sarcoma is characterized by its small-sized tumors with fibrovascular stroma sometimes with myxoid change. The tumor nuclei can be round or ovoid, but the common feature is the scant cytoplasm, like Ewing sarcoma. Although fusion proteins could theoretically be detected in the cytoplasm or ER, their minimal amount makes the interpretation of immunohistochemical stains challenging. The analysis of the disorder propensities by using IUPred3 revealed a significant increase in the disorder of BCOR^NTs^ in region^PUFD^ of the fusion proteins as compared to wBCOR. This increase may affect the interactions of BCOR with the PCGF1’s RAWUL domain and with the KDM2B’s C terminus, potentially interfering with the formation of the PRC1.1 complex.

We hypothesized that the fusion events may potentially have an impact on the interaction between the PUFD domain and the RAWUL domain of PCGF1. To test this hypothesis, we built the RAWUL-PUFD complexes and predicted their interaction energies and binding affinity. Ten different interaction types were analysed, and it was discovered that four of them - ICs charged-charged, ICs polar-polar, ICs charged-polar, and ICs charged-apolar - were not prominent in the fusion protein group as compared to the wild-type 4hpl. The fusion complexes were above the threshold for the ΔΔG, ICs polar-apolar, and NIS charged. However, only the BCOR::CCNB3, BCOR::MAML3, and RTL9::BCOR-2 fusions exceeded the threshold for apolar-apolar ICs. Additionally, the NIS apolar values remained unchanged except for BCOR::CCNB3 (**Fig. 4A**).

Based on the PRODIGY prediction, the binding affinity of the fusion complexes exhibited a decrease (**Fig. 4B,C**; **Table 5**; **Fig. 6**). To further investigate the binding interfaces, we expanded the repertoire of known interaction surfaces (**Fig. 5E**). Our findings suggest that the fusion event had an impact on the Interaction-Surface^NT/CT^ and the Interaction-SurfaceLeu (**Table 5**), as per the 0.05 cut-off value. Minor structural disparities were observed among the dimers (**Fig. 5C**). Our statistical analyses comparing the wild-type and fusion complexes revealed a significant change in the binding affinity of the studied structures (**Fig 6A**). Furthermore, this change was also revealed when specifically examining the Interaction surfaces (**Fig. 6B**), as well as when focusing solely on the Interaction-Surface^NT/CT^(**Fig. 6C**). Based on the analysis of IC and ΔG, the RAWUL-PUFD interactions demonstrated significant alterations and weakening, potentially impairing the assembly of PRC1.1.

The N terminus of BCOR contains a repression domain that interacts with a 60 kDa mitochondrial heat shock protein (HSPD1). This interaction is hypothesized to facilitate domain folding before chromatin association. The HSPD1-interacting region can independently inhibit transcription, which is distinct from the C terminus that plays a significant role in the recruitment to target sites. The linker region between ANK repeats and PUFD interacts with PCGF1, which recruits BCOR *via* direct interaction with KDM2B in wild-type hESCs. The BCOR-PRC1.1 complex is a suppressor of critical differentiation programs, highlighting BCOR’s central role in maintaining the pluripotent state and repressing mesoderm and endoderm specification (Wang *et al*, 2018). BCOR is involved in pluripotency and is a key regulator in various developmental processes, including embryogenesis, mesenchymal stem cell function, hematopoiesis, and lymphoid development. Disruptive mutations in BCOR and its paralog BCORL1 contribute to the pathogenesis of hematologic malignancies, exhibiting similarities to those observed in oculofaciocardiodental (OFCD) syndrome and Shukla-Vernon syndrome (Sportoletti *et al*, 2021). The BCOR K607E mutation near the Bbs has lesser affinities for BCL6, PCGF1, and RING1B proteins. Expression of the K607E mutant BCOR significantly increases cell proliferation, AKT phosphorylation, and interleukin-2 (IL-2) expression, with upregulated expression of HOX and S100 protein genes in T cells (Kang *et al*, 2021). Loss of BCOR function by deletion of its N-terminus (absence of Bbs) or C-terminus (absence of ANK and PUFD) is associated with reduced levels of H2AK119ub1 at PRC1 target promoters, including Hoxa7, Hoxa9, and Cebpa, highlighting its role in epigenetic regulation and suggesting its function as a tumor suppressor. BCOR mutations drive myelodysplastic syndromes and acute leukemia, especially in the presence of concurrent driver mutations, suggesting its function as a tumor suppressor (Tara *et al*, 2018).

Our findings posit that the functional integrity of BCOR fusion partner proteins is potentially compromised due to the loss of one or more functional domains, while the BCOR protein itself remains relatively intact across all instances. However, in the BCOR^NT^ group, the absence of Bbs may give rise to oncogenic effects, as evidenced in murine models (Tara *et al*, 2018). Notably, in the BCOR^CT^ group, despite the presence of Bbs, there is a conceivable attenuation in its interaction with BCL6, possibly attributed to physicochemical alterations. Furthermore, our results underscore the impact of protein fusions on the functionality of the PUFD domain. The observed alterations in interactions with the RAWUL domain suggest a potential disruption in the assembly of the PRC1.1. This perturbation in the complex formation may be attributed to the weakened binding affinity, thereby providing mechanistic insights into the oncogenic nature of *BCOR*-rearrangements in sarcomas. These nuanced molecular changes delineate the multifaceted nature of *BCOR* alterations and their consequential effects on protein functionality, intermolecular interactions, and the intricate regulatory networks governing cellular homeostasis. The elucidation of these mechanisms contributes to a more comprehensive understanding of the underlying oncogenic processes associated with *BCOR*-rearrangements in the context of sarcomas.

In conclusion, from the literature, we collected the chromosomal breakpoints and assembled the cDNA and protein sequences of seven known BRS. Our *in silico* analysis sheds light on the protein-level consequences of *BCOR* fusion events in small round cell sarcoma. These findings align with previous expression data indicating the upregulation of PRC1-regulated genes in BRS samples. The reduced binding affinity between RAWUL and PUFD domains, attributed to fusion events, may hinder dimer formation, and disrupt PRC1 assembly. The substantial alterations observed in both sequence and structural characteristics of the fusion proteins further underscore the oncogenic potential of BCOR-rearrangements and their contribution to the dysregulation of cellular processes in BRS. Overall, our study provides valuable insights into the oncogenic mechanism of BRS and underscores the potential of computational approaches in elucidating novel therapeutic targets for this rare soft tissue tumor.

## Methods

### Protein information

The nucleotide sequences of the seven fusion genes – that were determined by Sanger sequencing – were obtained from the scientific literature (Kao *et al*, 2018; Pierron *et al*, 2012; Specht *et al*, 2016; Yoshida *et al*, 2020; Vasella *et al*, 2022; Chang Chien *et al*, 2023). Ensemble (Genome assembly: GRCh37.p13; GCA_000001405.14) was used to find the breakpoints of the fusions (Martin *et al*, 2023) and to manually assemble the sequences. The sequences of the translated proteins were obtained based on the coding sequences by using the Expasy: Translate online tool (Gasteiger *et al*, 2003). Protein information for the BCOR, G2/mitotic-specific cyclin-B3 (CCNB3), mastermind-like protein 3 (MAML3), zinc finger CCCH domain-containing protein 7B (Z3H7B), histone-lysine N-methyltransferase 2D (KMT2D), MHC class II transactivator (CIITA), retrotransposon Gag-like protein 9 (RTL9), and calmegin (CLGN) proteins or their assembled fusion proteins were obtained from the UniProt database (The UniProt Consortium, 2023). The fusion of genes and proteins is indicated by a double colon (::) (Bruford *et al*, 2021).

The domain information was obtained from UniProt, the Conserved Domain database (CDD) (Lu *et al*, 2020), and InterPro (Paysan-Lafosse *et al*, 2023) databases. The Gene Ontology terms (GO) were determined based on the amino acid sequences we assembled on InterPro. The PANTHER GO terms were used (Ashburner *et al*, 2000; Mi *et al*, 2019a, 2019b; The Gene Ontology Consortium *et al*, 2023).

### Calculation of physicochemical properties

The physicochemical properties were calculated by using ExPASy’s ProtParam tool (Gasteiger *et al*, 2005), including the theoretical isoelectric point (TpI), molecular weight, the total number of positive and negatively charged residues, instability index (II) (Guruprasad *et al*, 1990), aliphatic index (AI) (Ikai, 1980), and grand average hydrophobicity (GRAVY) (Kyte & Doolittle, 1982). The instability index provides an estimate of the *in vitro* stability of a protein. The aliphatic index of a protein is considered a positive factor in increasing the thermostability of globular proteins and is specifically defined as the relative volume occupied by aliphatic side chains (alanine, valine, isoleucine, and leucine). The GRAVY score is calculated as the sum of the hydropathy values of all amino acids (AA) divided by the number of residues in the sequence. The following formula was used to calculate the change in physicochemical values between the wild-type partner and the fusion proteins:

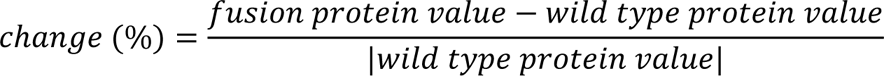

### Prediction of signal peptides and disorder propensities

The SignalP 6.0 predictor was utilized to identify signal peptides in the proteins. Only Sec/Signal Peptides (Sec/SPI) found in eukaryotes were searched, as the selected input organism was Eukarya (Teufel *et al*, 2022). DeepLoc 2.0 was utilized to determine the intracellular localization of the proteins. Only the reported probability scores above the thresholds set by DeepLoc 2.0 were considered: cytoplasm (0.4761), nucleus (0.5014), and endoplasmic reticulum (0.6090) (Thumuluri *et al*, 2022). To validate the predictor’s results, we also utilized the UniProt database and The Human Protein Atlas (Thul *et al*, 2017; The UniProt Consortium, 2023). Additionally, the NetGPI-1.1 GPI-anchored predictor (Gíslason *et al*, 2021) was applied to determine the potential glycosylphosphatidylinositol (GPI) anchoring of the proteins.

We used the IUPred3 web server to compare the 1448-1633 (region^ANK+linker^) and 1634-1748 (region^PUFD^) regions of the wild-type BCOR (wBCOR) with its representative intrinsically disordered regions (IDRs) in the fusion proteins (Erdős *et al*, 2021).

### Analysis of protein structures

AlphaFold2 and AlphaFold2-multimer (AF2, version 2.3.2) were utilized to predict the 3D structures of the fusion protein complexes. The predictions resulted in five structures per prediction, ranked by AF2. The top-ranked models were selected for in-depth investigation of intermolecular contacts (ICs), binding affinity (ΔG) prediction, and contact map generation. Meanwhile, all five structures per predicted protein were used for statistical comparison, ensuring a comprehensive analysis of predictive performance across multiple structural models. The structure of KMT2D::BCOR could not be predicted due to the length of the sequence (5480 AA) that exceeds the limit of the computational algorithm. AF2 was accessed through the COSMIC2 Science Gateway (Evans *et al*, 2022; Cianfrocco *et al*, 2017; Tunyasuvunakool *et al*, 2021).

The PRODIGY web server was used to predict the binding affinity described through, Gibbs free energy ΔG (kcal mol^-1^), and ICs between the chains of the protein complexes. The smaller the Kd and the ΔG value, the greater the binding affinity. To compare the predicted structures, we used an experimentally determined structure of BCOR (PDB ID: 4hpl) (Junco *et al*, 2013) that was downloaded from the RCSB Protein Data Bank (Burley *et al*, 2023). We generated the quaternary structures of the fusion proteins complexed with PCGF1 by AF2. To ensure comparability with the chosen control, and to limit PRODIGY predictions to the RAWUL-PUFD sequences relevant to our study, we studied the complexes containing the 167-177 and 185-254 regions of the RAWUL domain as well as the 1636-1748 region of the PUFD domain, resembling the regions that are included in the 4hpl.pdb coordinate file. To visualize the protein structures, PyMOL was used. (The PyMOL Molecular Graphics System, Version 2.5, Schrödinger, LLC). The loop encompassing the 178-184 residues of the RAWUL domain is missing from the crystal structure (4hpl), therefore, this region was omitted from the analyses. Although this region is not part of the interface between the RAWUL and PUFD domains, it was removed from the modeled complexes, as well, by using PyMOL (The PyMOL Molecular Graphics System, Version 2.5, Schrödinger, LLC). For comparison, a modeled complex was also prepared based on the crystal structure (4hpl) by using AF2 (4hpl^AF2^).

To identify major changes among rank1 models in intermolecular contacts (ICs) resulting from the fusion event in the novel protein dimers, a predetermined cut-off value was employed. This cut-off was determined by comparing IC values between the experimental structure (4hpl) and the predicted control (4hpl^AF2^). Specifically, deviations exceeding the difference between IC values at 4hpl and 4hpl^AF2^ were considered noteworthy. Evaluation of ICs in the RAWUL-PUFD dimers within the fusion proteins was conducted relative to the IC values observed in the 4hpl structure, with a deviation of 2 ICs from these values considered indicative of meaningful changes. Additionally, thresholds for changes in the NIS charged and NIS apolar parameters were established at differences of 0.16% and 0.25%, respectively (**Fig. 4a**). This systematic approach ensures a meticulous assessment of alterations in intermolecular contacts, utilizing well-defined thresholds derived from control values.

To calculate the change in binding affinity (ΔΔG) between wild-type and fusion-type complexes resulting from fusion events, we calculated ΔΔG using the following formula:

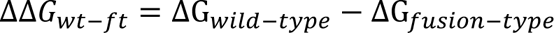

To interpret ΔΔG results values less than zero indicate a decrease in binding affinity between the RAWUL and PUFD domains, while values greater than zero indicate an increase. In our study, to compare the binding affinity values between wild-type (4hpl) and fusion-type (BCOR fusion proteins) proteins and to distinguish notable ΔΔG values, we encountered a lack of established literature on the subject. Therefore, we adopted a pragmatic approach and determined a cut-off of 0.5 for distinguishing major increases and decreases in ΔΔG values. This cut-off was derived like the commonly used cut-off for ΔΔG values in the context of single nucleotide polymorphism mutations, as described in the literature(Khan & Vihinen, 2010). This threshold defines ΔΔG ≥ 0.5 as a relevant increase or ΔΔG ≤ -0.5 as a relevant decrease in binding affinity, providing a quantitative criterion for evaluating the impact of structural modifications on protein-protein interactions. The contact maps were generated using MAPIYA with a distance cut-off of 5.5 Å. Possible interaction forces were also described in brackets after the contacts (Badaczewska-Dawid *et al*, 2022).

### Statistics

A non-parametric Friedman’s test was performed to compare the IUPred3 IDRs scores of wBCOR and the IDRs scores of fusion proteins. As post hoc, Dunn’s multiple comparison tests were performed to compare each of the fusion proteins to the wBCOR.

Friedman’s tests followed by Dunn’s multiple comparison tests were performed, using GraphPad Prism version 9.5.1 (Friedman, 1937; Dunn, 1964; GraphPad Prism version 9.5.1 for Windows, GraphPad Software, Boston, Massachusetts USA, www.graphpad.com).

Ordinary one-way ANOVA and Tukey’s multiple comparison tests were used to compare the binding affinity values of the complexes. *P* values were considered significant where *P* < 0.05.

For graph and figure construction, GraphPad Prism (GraphPad Prism version 9.5.1 for Windows, GraphPad Software, Boston, Massachusetts USA, www.graphpad.com) version 9.5.1 and IBS (illustrator of biological sequences) were used (Liu *et al*, 2015).

## Acknowledgements

Kristóf Madarász was supported by PhD Excellence Scholarship from Count István Tisza Foundation for the University of Debrecen and by the ÚNKP-23-3-II-DE-330 New National Excellence Program of the Ministry of Innovation and Technology.

This project was supported by the ÚNKP-23-5-DE-486 New National Excellence Program of the Ministry of Culture and Innovation from the source of the National Research, Development and Innovation Fund (to J.AM.). János A. Mótyán is the receiver of the János Bolyai Research Scholarship provided by the Hungarian Academy of Sciences (BO/00110/23/5).

## Authors’ contributions

K.M. collected and analysed the data, prepared the figures, designed, and wrote the manuscript. J.A.M. supervised, revised, and wrote the manuscript. Y.-C.C.C. provided expertise in the field of sarcoma. J.B. and S.L.C. provided valuable suggestions. G.M. supervised the project, provided the resources, revised and edited the final manuscript. A.M. conceptualized, supervised, and coordinated the project, wrote, revised, and edited the manuscript. All authors read and approved the final manuscript.

## Conflict of interest

The authors declare that they have no competing interests.

## Data Availability Section

The datasets used and/or analysed during the current study are available from the corresponding author on reasonable request. This study includes no data deposited in external repositories.

## References

Albagli O, Lantoine D, Quief S, Quignon F, Englert C, Kerckaert J-P, Montarras D, Pinset C & Lindon C (1999) Overexpressed BCL6 (LAZ3) oncoprotein triggers apoptosis, delays S phase progression and associates with replication foci. Oncogene 18: 5063–5075

Ashburner M, Ball CA, Blake JA, Botstein D, Butler H, Cherry JM, Davis AP, Dolinski K, Dwight SS, Eppig JT, et al (2000) Gene Ontology: tool for the unification of biology. Nat Genet 25: 25–29

Badaczewska-Dawid AE, Nithin C, Wroblewski K, Kurcinski M & Kmiecik S (2022) MAPIYA contact map server for identification and visualization of molecular interactions in proteins and biological complexes. Nucleic Acids Research 50: W474–W482

Blackledge NP, Farcas AM, Kondo T, King HW, McGouran JF, Hanssen LLP, Ito S, Cooper S, Kondo K, Koseki Y, et al (2014) Variant PRC1 Complex-Dependent H2A Ubiquitylation Drives PRC2 Recruitment and Polycomb Domain Formation. Cell 157: 1445–1459

Blackledge NP, Rose NR & Klose RJ (2015) Targeting Polycomb systems to regulate gene expression: modifications to a complex story. Nat Rev Mol Cell Biol 16: 643–649

Brown RS (2005) Zinc finger proteins: getting a grip on RNA. Curr Opin Struct Biol 15: 94–98

Bruford EA, Antonescu CR, Carroll AJ, Chinnaiyan A, Cree IA, Cross NCP, Dalgleish R, Gale RP, Harrison CJ, Hastings RJ, et al (2021) HUGO Gene Nomenclature Committee (HGNC) recommendations for the designation of gene fusions. Leukemia 35: 3040–3043

Burley SK, Bhikadiya C, Bi C, Bittrich S, Chao H, Chen L, Craig PA, Crichlow GV, Dalenberg K, Duarte JM, et al (2023) RCSB Protein Data Bank (RCSB.org): delivery of experimentally-determined PDB structures alongside one million computed structure models of proteins from artificial intelligence/machine learning. Nucleic Acids Research 51: D488–D508

Bushweller JH, Schmidt C, Achille N, Kuntimaddi A, Boulton A, Leach B, Zhang S & Zeleznik-Le NJ (2018) Direct Binding of BCOR, but Not CBX8, to MLL-AF9 Is Essential for MLL-AF9 Leukemia Via Regulation of the EYA1/SIX1 Gene Network. Blood 132: 1316

Campillos M, Doerks T, Shah PK & Bork P (2006) Computational characterization of multiple Gag-like human proteins. Trends in Genetics 22: 585–589

Carballo E, Lai WS & Blackshear PJ (1998) Feedback inhibition of macrophage tumor necrosis factor-alpha production by tristetraprolin. Science 281: 1001–1005

Chang Chien Y-C, Madarász K, Csoma SL, Mótyán JA, Huang H-Y, Méhes G & Mokánszki A (2023) Molecular Identification and In Silico Protein Analysis of a Novel BCOR-CLGN Gene Fusion in Intrathoracic BCOR-Rearranged Sarcoma. Cancers 15: 898

Chiang MY, Xu ML, Histen G, Shestova O, Roy M, Nam Y, Blacklow SC, Sacks DB, Pear WS & Aster JC (2006) Identification of a conserved negative regulatory sequence that influences the leukemogenic activity of NOTCH1. Mol Cell Biol 26: 6261–6271

Chittock EC, Latwiel S, Miller TCR & Müller CW (2017) Molecular architecture of polycomb repressive complexes. Biochemical Society Transactions 45: 193–205

Cho Y-W, Hong T, Hong S, Guo H, Yu H, Kim D, Guszczynski T, Dressler GR, Copeland TD, Kalkum M, et al (2007) PTIP Associates with MLL3-and MLL4-containing Histone H3 Lysine 4 Methyltransferase Complex*♦. Journal of Biological Chemistry 282: 20395–20406

Cianfrocco MA, Wong-Barnum M, Youn C, Wagner R & Leschziner A (2017) COSMIC2: A Science Gateway for Cryo-Electron Microscopy Structure Determination. In Proceedings of the Practice and Experience in Advanced Research Computing 2017 on Sustainability, Success and Impact pp 1–5. New York, NY, USA: Association for Computing Machinery

Dunn OJ (1964) Multiple Comparisons Using Rank Sums. Technometrics 6: 241–252

Erdős G, Pajkos M & Dosztányi Z (2021) IUPred3: prediction of protein disorder enhanced with unambiguous experimental annotation and visualization of evolutionary conservation. Nucleic Acids Research 49: W297–W303

Evans R, O’Neill M, Pritzel A, Antropova N, Senior A, Green T, Žídek A, Bates R, Blackwell S, Yim J, et al (2022) Protein complex prediction with AlphaFold-Multimer. 2021.10.04.463034 doi:10.1101/2021.10.04.463034 [PREPRINT]

Friedman M (1937) The Use of Ranks to Avoid the Assumption of Normality Implicit in the Analysis of Variance. Journal of the American Statistical Association 32: 675–701

Gasteiger E, Gattiker A, Hoogland C, Ivanyi I, Appel RD & Bairoch A (2003) ExPASy: the proteomics server for in-depth protein knowledge and analysis. Nucleic Acids Res 31: 3784– 3788

Gasteiger E, Hoogland C, Gattiker A, Duvaud S, Wilkins MR, Appel RD & Bairoch A (2005) Protein Identification and Analysis Tools on the ExPASy Server. In The Proteomics Protocols Handbook, Walker JM (ed) pp 571–607. Totowa, NJ: Humana Press

Ghetu AF, Corcoran CM, Cerchietti L, Bardwell VJ, Melnick A & Privé GG (2008) Structure of a BCOR corepressor peptide in complex with the BCL6 BTB domain dimer. Mol Cell 29: 384– 391

Gíslason MH, Nielsen H, Almagro Armenteros JJ & Johansen AR (2021) Prediction of GPI-anchored proteins with pointer neural networks. Current Research in Biotechnology 3: 6–13

GraphPad Prism version 9.5.1 for Windows, GraphPad Software, Boston, Massachusetts USA, www.graphpad.com

Guruprasad K, Reddy BV & Pandit MW (1990) Correlation between stability of a protein and its dipeptide composition: a novel approach for predicting in vivo stability of a protein from its primary sequence. Protein Eng 4: 155–161

Huynh KD, Fischle W, Verdin E & Bardwell VJ (2000) BCoR, a novel corepressor involved in BCL-6 repression. Genes Dev 14: 1810–1823

Ikai A (1980) Thermostability and aliphatic index of globular proteins. J Biochem 88: 1895–1898

Ikawa M, Nakanishi T, Yamada S, Wada I, Kominami K, Tanaka H, Nozaki M, Nishimune Y & Okabe M (2001) Calmegin Is Required for Fertilin α/β Heterodimerization and Sperm Fertility. Developmental Biology 240: 254–261

Ikawa M, Wada I, Kominami K, Watanabe D, Toshimori K, Nishimune Y & Okabe M (1997) The putative chaperone calmegin is required for sperm fertility. Nature 387: 607–611

Ishino F, Itoh J, Irie M, Matsuzawa A, Naruse M, Suzuki T, Hiraoka Y & Kaneko-Ishino T (2023) Retrovirus-Derived RTL9 Plays an Important Role in Innate Antifungal Immunity in the Eutherian Brain. Int J Mol Sci 24: 14884

Junco SE, Wang R, Gaipa JC, Taylor AB, Schirf V, Gearhart MD, Bardwell VJ, Demeler B, Hart PJ & Kim CA (2013) Structure of the Polycomb Group Protein PCGF1 in Complex with BCOR Reveals Basis for Binding Selectivity of PCGF Homologs. Structure 21: 665–671

Kang JH, Lee SH, Lee J, Choi M, Cho J, Kim SJ, Kim WS, Ko YH & Yoo HY (2021) The mutation of BCOR is highly recurrent and oncogenic in mature T-cell lymphoma. BMC Cancer 21: 82

Kao Y-C, Owosho AA, Sung Y-S, Zhang L, Fujisawa Y, Lee J-C, Wexler L, Argani P, Swanson D, Dickson BC, et al (2018) BCOR-CCNB3-Fusion Positive Sarcomas: A Clinicopathologic and Molecular Analysis of 36 cases with Comparison to Morphologic Spectrum and Clinical Behavior of other Round Cell Sarcomas. Am J Surg Pathol 42: 604–615

Kao Y-C, Sung Y-S, Zhang L, Huang S-C, Argani P, Chung CT, Graf NS, Wright DC, Kellie SJ, Agaram NP, et al (2016) Recurrent BCOR Internal Tandem Duplication and YWHAE-NUTM2B Fusions in Soft Tissue Undifferentiated Round Cell Sarcoma of Infancy: Overlapping Genetic Features With Clear Cell Sarcoma of Kidney. Am J Surg Pathol 40: 1009–1020

Khan S & Vihinen M (2010) Performance of protein stability predictors. Human Mutation 31: 675– 684

Kyte J & Doolittle RF (1982) A simple method for displaying the hydropathic character of a protein. J Mol Biol 157: 105–132

Lai WS, Carballo E, Strum JR, Kennington EA, Phillips RS & Blackshear PJ (1999) Evidence that tristetraprolin binds to AU-rich elements and promotes the deadenylation and destabilization of tumor necrosis factor alpha mRNA. Mol Cell Biol 19: 4311–4323

Lee HS, Bang I, You J, Jeong T-K, Kim CR, Hwang M, Kim J-S, Baek SH, Song J-J & Choi H-J (2023) Molecular basis for PHF7-mediated ubiquitination of histone H3. Genes Dev 37: 984– 997

León Machado JA & Steimle V (2021) The MHC Class II Transactivator CIITA: Not (Quite) the Odd-One-Out Anymore among NLR Proteins. International Journal of Molecular Sciences 22: 1074

Linhoff MW, Harton JA, Cressman DE, Martin BK & Ting JP (2001) Two distinct domains within CIITA mediate self-association: involvement of the GTP-binding and leucine-rich repeat domains. Mol Cell Biol 21: 3001–3011

Liu H, Kennard S & Lilly B (2009) NOTCH3 expression is induced in mural cells through an autoregulatory loop that requires endothelial-expressed JAGGED1. Circ Res 104: 466–475

Liu W, Xie Y, Ma J, Luo X, Nie P, Zuo Z, Lahrmann U, Zhao Q, Zheng Y, Zhao Y, et al (2015) IBS: an illustrator for the presentation and visualization of biological sequences. Bioinformatics 31: 3359–3361

Lu S, Wang J, Chitsaz F, Derbyshire MK, Geer RC, Gonzales NR, Gwadz M, Hurwitz DI, Marchler GH, Song JS, et al (2020) CDD/SPARCLE: the conserved domain database in 2020. Nucleic Acids Res 48: D265–D268

Martin FJ, Amode MR, Aneja A, Austine-Orimoloye O, Azov AG, Barnes I, Becker A, Bennett R, Berry A, Bhai J, et al (2023) Ensembl 2023. Nucleic Acids Research 51: D933–D941

Mi H, Muruganujan A, Ebert D, Huang X & Thomas PD (2019a) PANTHER version 14: more genomes, a new PANTHER GO-slim and improvements in enrichment analysis tools. Nucleic Acids Research 47: D419–D426

Mi H, Muruganujan A, Huang X, Ebert D, Mills C, Guo X & Thomas PD (2019b) Protocol Update for large-scale genome and gene function analysis with the PANTHER classification system (v.14.0). Nat Protoc 14: 703–721

Nam Y, Sliz P, Song L, Aster JC & Blacklow SC (2006) Structural basis for cooperativity in recruitment of MAML coactivators to Notch transcription complexes. Cell 124: 973–983

Nguyen TB, Manova K, Capodieci P, Lindon C, Bottega S, Wang X-Y, Refik-Rogers J, Pines J, Wolgemuth DJ & Koff A (2002) Characterization and Expression of Mammalian Cyclin B3, a Prepachytene Meiotic Cyclin *. Journal of Biological Chemistry 277: 41960–41969

Pagan JK, Arnold J, Hanchard KJ, Kumar R, Bruno T, Jones MJK, Richard DJ, Forrest A, Spurdle A, Verdin E, et al (2007) A Novel Corepressor, BCoR-L1, Represses Transcription through an Interaction with CtBP. Journal of Biological Chemistry 282: 15248–15257

Paysan-Lafosse T, Blum M, Chuguransky S, Grego T, Pinto BL, Salazar GA, Bileschi ML, Bork P, Bridge A, Colwell L, et al (2023) InterPro in 2022. Nucleic Acids Research 51: D418–D427

Pierron G, Tirode F, Lucchesi C, Reynaud S, Ballet S, Cohen-Gogo S, Perrin V, Coindre J-M & Delattre O (2012) A new subtype of bone sarcoma defined by BCOR-CCNB3 gene fusion. Nat Genet 44: 461–466

Sbaraglia M, Bellan E & Dei Tos AP (2020) The 2020 WHO Classification of Soft Tissue Tumours: news and perspectives. Pathologica 113: 70–84

Schwartz R, Ting CS & King J (2001) Whole Proteome pI Values Correlate with Subcellular Localizations of Proteins for Organisms within the Three Domains of Life. Genome Res 11: 703–709

Segel M, Lash B, Song J, Ladha A, Liu CC, Jin X, Mekhedov SL, Macrae RK, Koonin EV & Zhang F (2021) Mammalian retrovirus-like protein PEG10 packages its own mRNA and can be pseudotyped for mRNA delivery. Science 373: 882–889

Specht K, Zhang L, Sung Y-S, Nucci M, Dry S, Vaiyapuri S, Richter GHS, Fletcher CDM & Antonescu CR (2016) Novel BCOR-MAML3 and ZC3H7B-BCOR Gene Fusions in Undifferentiated Small Blue Round Cell Sarcomas. Am J Surg Pathol 40: 433–442

Sportoletti P, Sorcini D & Falini B (2021) BCOR gene alterations in hematologic diseases. Blood 138: 2455–2468

Srinivasan RS, Erkenez AC de & Hemenway CS (2003) The mixed lineage leukemia fusion partner AF9 binds specific isoforms of the BCL-6 corepressor. Oncogene 22: 3395–3406

Srivastava A, Nagai T, Srivastava A, Miyashita O & Tama F (2018) Role of Computational Methods in Going beyond X-ray Crystallography to Explore Protein Structure and Dynamics. Int J Mol Sci 19: 3401

Tara S, Isshiki Y, Nakajima-Takagi Y, Oshima M, Aoyama K, Tanaka T, Shinoda D, Koide S, Saraya A, Miyagi S, et al (2018) Bcor insufficiency promotes initiation and progression of myelodysplastic syndrome. Blood 132: 2470–2483

Teufel F, Almagro Armenteros JJ, Johansen AR, Gíslason MH, Pihl SI, Tsirigos KD, Winther O, Brunak S, von Heijne G & Nielsen H (2022) SignalP 6.0 predicts all five types of signal peptides using protein language models. Nat Biotechnol 40: 1023–1025

The Gene Ontology Consortium, Aleksander SA, Balhoff J, Carbon S, Cherry JM, Drabkin HJ, Ebert D, Feuermann M, Gaudet P, Harris NL, et al (2023) The Gene Ontology knowledgebase in 2023. Genetics 224: iyad031

The PyMOL Molecular Graphics System, Version 2.5, Schrödinger, LLC

The UniProt Consortium (2023) UniProt: the Universal Protein Knowledgebase in 2023. Nucleic Acids Research 51: D523–D531

Thul PJ, Åkesson L, Wiking M, Mahdessian D, Geladaki A, Ait Blal H, Alm T, Asplund A, Björk L, Breckels LM, et al (2017) A subcellular map of the human proteome. Science 356: eaal3321

Thumuluri V, Almagro Armenteros JJ, Johansen AR, Nielsen H & Winther O (2022) DeepLoc 2.0: multi-label subcellular localization prediction using protein language models. Nucleic Acids Research 50: W228–W234

Treiber T, Treiber N, Plessmann U, Harlander S, Daiß J-L, Eichner N, Lehmann G, Schall K, Urlaub H & Meister G (2017) A Compendium of RNA-Binding Proteins that Regulate MicroRNA Biogenesis. Molecular Cell 66: 270–284.e13

Tunyasuvunakool K, Adler J, Wu Z, Green T, Zielinski M, Žídek A, Bridgland A, Cowie A, Meyer C, Laydon A, et al (2021) Highly accurate protein structure prediction for the human proteome. Nature 596: 590–596

Vasella M, Wagner U, Fritz C, Seidl K, Giudici L, Exner GU, Moch H, Wild PJ & Bode-Lesniewska B (2022) Novel RGAG1-BCOR gene fusion revealed in a somatic soft tissue sarcoma with a long follow-up. Virchows Arch 480: 1107–1114

Wamstad JA, Corcoran CM, Keating AM & Bardwell VJ (2008) Role of the Transcriptional Corepressor Bcor in Embryonic Stem Cell Differentiation and Early Embryonic Development. PLoS ONE 3: e2814

Wang Z, Gearhart MD, Lee Y-W, Kumar I, Ramazanov B, Zhang Y, Hernandez C, Lu AY, Neuenkirchen N, Deng J, et al (2018) A non-canonical BCOR-PRC1.1 complex represses differentiation programs in human ESCs. Cell Stem Cell 22: 235–251.e9

Watanabe D, Yamada K, Nishina Y, Tajima Y, Koshimizu U, Nagata A & Nishimune Y (1994) Molecular cloning of a novel Ca(2+)-binding protein (calmegin) specifically expressed during male meiotic germ cell development. Journal of Biological Chemistry 269: 7744–7749

Wong SJ, Senkovich O, Artigas JA, Gearhart MD, Ilangovan U, Graham DW, Abel KN, Yu T, Hinck AP, Bardwell VJ, et al (2020) Structure and Role of BCOR PUFD in Noncanonical PRC1 Assembly and Disease. Biochemistry 59: 2718–2728

Wu L, Maillard I, Nakamura M, Pear WS & Griffin JD (2007) The transcriptional coactivator Maml1 is required for Notch2-mediated marginal zone B-cell development. Blood 110: 3618–3623

Yoshida A, Arai Y, Hama N, Chikuta H, Bando Y, Nakano S, Kobayashi E, Shibahara J, Fukuhara H, Komiyama M, et al (2020) Expanding the clinicopathologic and molecular spectrum of BCOR-associated sarcomas in adults. Histopathology 76: 509–520

